# Functional modules within a distributed neural network control feeding in a model medusa

**DOI:** 10.1101/2021.02.22.432372

**Authors:** Brandon Weissbourd, Tsuyoshi Momose, Aditya Nair, Ann Kennedy, Bridgett Hunt, David J. Anderson

**Affiliations:** Division of Biology and Biological Engineering; Howard Hughes Medical Institute; Tianqiao and Chrissy Chen Institute for Neuroscience California Institute of Technology, Pasadena, CA 91125, USA; Sorbonne Université, CNRS, Laboratoire de Biologie du Développement de Villefranche-sur-mer (LBDV), 06230 Villefranche-sur-mer, France; Department of Physiology, Feinberg School of Medicine Northwestern University, Chicago, IL 60611, USA

## Abstract

Jellyfish are free-swimming, radially symmetric organisms with complex behaviors that arise from coordinated interactions between distinct, autonomously functioning body parts. This behavioral complexity evolved without a corresponding cephalization of the nervous system. The systems-level neural mechanisms through which such decentralized control is achieved remain unclear. Here, we address this question using the jellyfish, *Clytia,* and present it as a new neuroscience model. We describe a coordinated, asymmetric behavior in which food is passed from the umbrellar margin to the central mouth via directed margin folding. Using newly developed transgenic jellyfish lines to ablate or image specific neuronal subpopulations, we find, unexpectedly, that margin folding reflects the local activation of neural subnetworks that tile the umbrella. Modeling suggests that this structured ensemble activity emerges from sparse, local connectivity rules. These findings reveal how an organismal behavior can emerge from local interactions between functional modules in the absence of a central brain.

## Introduction

Neurons first appeared in evolution prior to the last common ancestor of cnidarians (a phylum of radially symmetric invertebrates) and bilaterians (Figure 1A; Bucher and Anderson, 2015; Arendt et al., 2016; Kristan, 2016). From this simple beginning, there has been massive diversification of nervous system structure and function in both cnidarian and bilaterian lineages, as evidenced by the incredible diversity of animal behaviors and sensory-motor capabilities. However, whether this behavioral diversity reflects a similar diversity of underlying neural mechanisms, or if there are principles that are conserved or converged upon, remains unclear.

**Figure 1:**
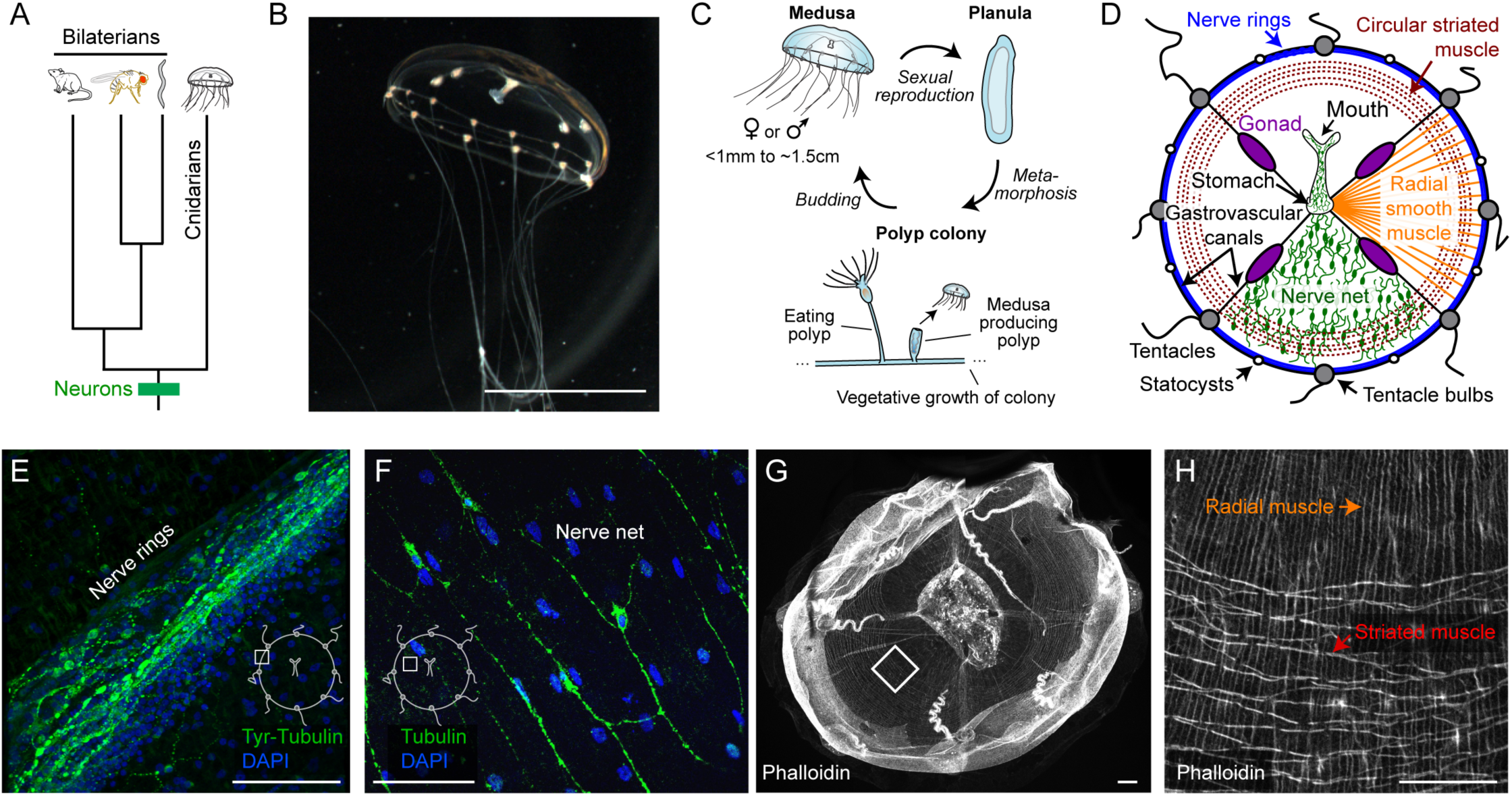
Introduction to *Clytia*. (A) Phylogenetic relationship between cnidarians and several bilaterian lineages. Neurons appeared prior to their last common ancestor. (B) A *Clytia* medusa. Orange coloration derives from food (brine shrimp). Scale ∼0.5cm. (C) *Clytia* life cycle. Medusae (jellyfish) are male or female; fertilized eggs develop into larvae (planulae) that metamorphose into primary polyps, which then grow into polyp colonies. A colony is composed of two types of polyps: one that eats, and one that releases baby jellyfish. Modified from Leclère and Röttinger, 2017. (D) Diagram of medusa anatomy. The nerve net and radial muscle are distributed across the animal but are shown in only one quadrant here for clarity. Modified from Houliston et al., 2010. (E) The nerve rings. Green: anti-Tyrosine Tubulin. Blue: DAPI. Scale: 50µm. Inset diagram here and in panel F indicates anatomical location. (F) The nerve net. Green: anti-*a*Tubulin. Blue: DAPI. Scale: 50µm. (G) Musculature in a whole animal (phalloidin staining). Scale: 100µm. (H) Magnified view of the boxed region in panel G showing the radial versus striated, circular muscle. Scale: 50µm.

Perhaps the most fundamental divergence in nervous system structure and function is the invention of cephalization in the bilaterian lineage: the coordinated control of behavior by a central ganglion (“brain”) located in the head (Arendt et al., 2008; Holland, 2003). While cnidarians exhibit behaviors requiring coordination between multiple body parts, their nervous systems lack such centralization. Instead, cnidarian nervous systems are distributed across the body (Lin et al., 2001; Satterlie, 2011; Burnett and Diehl, 1964; Bosch et al., 2017). Yet such organisms are able to feed themselves, reproduce sexually, escape from predators, and even sleep (Mackie, 2003, 2004, 1990, 1984; Fossette et al., 2015; Nath et al., 2017; Lewis and Long, 2005; Robson, 1961; Garm et al., 2007). How the neural control of such coordinated behaviors and internal states evolved in the absence of centralization remains unclear.

Among cnidarians, the greatest behavioral complexity is exhibited by the free-swimming life stages of medusozoans, otherwise known as jellyfish (Costello et al., 2008; Mackie, 2004, 1990; Satterlie, 2002; Costello et al., 2021). This behavioral complexity arises from deeply branching, independently evolved body configurations composed of multiple, distinct organs (Cartwright et al., 2007; Gold et al., 2019; Khalturin et al., 2019). Jellyfish are thought to have arisen more than 500 million years ago, and have since diversified into several thousand known species, divided into several classes (Khalturin et al., 2019; Collins et al., 2006; Zapata et al., 2015; Bouillon and Boero, 2000; Kramp, 1961; Liebeskind et al., 2016; catalogueoflife.org). Jellyfish often have complex lifecycles consisting of both motile and sessile (polyp colony) stages (Figure 1C; Leclère et al., 2019). In recent years, jellyfish have gained increasing interest as critical components of ocean ecosystems, in part due to the changing frequency of jellyfish blooms and their negative impact on commercial fishing, aquaculture, tourism, and even coastal power plants (Condon et al., 2013; Graham et al., 2014; Hays et al., 2018; Lynam et al., 2006). Yet remarkably little is known about the neural control of behavior in these organisms, especially at the systems level.

The decentralized nature of the jellyfish nervous system is most pronounced in the class *Hydrozoa* (Passano, 1973, 1965; Romanes, 1885; Satterlie, 2002). Hydrozoan nervous systems have modifications on a common organizational theme, in which most of the nervous system is arranged into concentric rings of neurons at the margin of the bell, or umbrella (“nerve rings”; Figure 1D; Hertwig and Hertwig, 1878; Satterlie and Spencer, 1983; Koizumi et al., 2015; Jha and Mackie, 1967; Anderson and Mackie, 1977; Mackie, 2004). Motor control of the umbrella (excluding mouth, tentacles, and velum) comes from two types of muscle: radially oriented smooth muscle, and circularly oriented, striated muscle (Mackie and Passano, 1968; Satterlie, 2008; Leclère and Röttinger, 2017). The remainder of the nervous system is distributed across the tentacles, gastrovascular canals, mouth, and in the subumbrella (Figure 1D; Lin et al., 2001; Mackie et al., 1985; Mackie and Meech, 2008). Neurons of the subumbrella often form diffuse “nerve nets” (Satterlie, 2008; Satterlie and Spencer, 1983; Mackie and Meech, 2008; Satterlie, 2015). However, jellyfish are capable of oriented, asymmetric movements of the umbrella, such as in feeding (Hyman, 1940; Satterlie, 2008; Spencer, 1975). How such vectorial behaviors are controlled at the systems level is unknown.

Jellyfish nervous systems have been studied extensively using single-unit electrophysiological recordings, but this approach makes it difficult to study neural systems-level properties. Breakthroughs in genetic and optical techniques now present the opportunity to study the function of such a nervous system *in toto* (Ahrens and Engert, 2015). Here we introduce the hydrozoan jellyfish *Clytia hemisphaerica,* originally developed to study early development and evolution (Houliston et al., 2010), as a new neuroscience model. This species met key criteria for selection: 1) small, transparent, and quasi-planar, facilitating optical approaches; 2) genetic manipulability; 3) a relatively numerically simple nervous system; 4) a suite of coordinated organismal behaviors; and, 5) adaptability to the laboratory, including ease of sexual reproduction. The *Clytia* genome has been sequenced (Leclère et al., 2019) and an atlas of its cell types generated using single-cell RNA sequencing (scRNAseq; Chari et al., 2021). CRISPR/Cas9-mediated gene knockout has also been achieved in this species (Momose et al., 2018).

In this inaugural study, we develop transgenesis in *Clytia* for the ablation and imaging of specific neuronal subpopulations. Using these tools, we have investigated the role of one such subpopulation in the control of an asymmetric, oriented feeding behavior that requires coordination between distinct body parts, each of which can perform a component action of the complete behavior in isolation. Using population imaging and modeling, we discover an unanticipated functional parcellation of the subumbrellar neural network that may explain such asymmetric behavior, contravening a prevailing view of this system as diffuse and unstructured. This work establishes *Clytia* as a genetically and physiologically tractable neuroscience model, and reveals unexpected organizational properties of its nervous system. This provides an entry point to investigate both the particulars of a jellyfish neural system, including how coordinated behavior is achieved in the absence of a central brain, as well as principles of neuroscience across phylogeny and levels of analysis.

## Results

*Clytia* medusae (jellyfish) are ∼1mm-1.5cm in diameter, radially symmetric, and optically transparent, with approximately 10,000 neurons in a 1cm animal (Figure 1B; Chari et al., 2021). Importantly, the complete, tri-phasic life cycle can be cultured under laboratory conditions (Figure 1C; Houliston et al., 2010; Lechable et al., 2020). Briefly, *Clytia* medusae have separate sexes that spawn daily. Embryos develop overnight into planula larvae, which, two days later, can be induced to metamorphose into a polyp attached to a microscope slide. It will then grow along this slide into a polyp colony, which is vegetatively propagating and considered immortal. After several weeks, the colony will begin to release jellyfish. A single polyp slide can produce dozens of clonal jellyfish overnight, which reach sexual maturity in 2-3 weeks. Our histological analysis in *Clytia* illustrates the hallmarks of the hydrozoan body plan described above (Figure 1D): nerve rings (Figure 1E), subumbrellar nerve net (Figure 1F), and both muscle types (Figure 1G-H).

### Margin folding is a coordinated, directionally targeted behavior arising from body parts that perform subfunctions autonomously

One of the most prominent coordinated behaviors exhibited by *Clytia* medusae is a margin folding action essential to feeding (Figure 2A; Hyman, 1940; Miglietta et al., 2000; Romanes, 1885; Satterlie, 1985; Spencer, 1975). Following prey capture by a tentacle, the tentacle retracts, swimming stops (Figure S1A), and the margin folds inwards to pass the food directionally to the mouth (Figure 2A, Supplemental Video 1). The mouth is located at the tip of an elongated feeding organ suspended from the roof of the umbrella (Figure 1B, D). This feeding organ (“mouth”) bends and orients towards the folding portion of the margin to facilitate passing of the prey item (“pointing”, Figure 2B). Food passing is robust and reproducible: 96% of first passing attempts occurred within one minute of prey (brine shrimp) capture, and of these 88% were successful (Figure S1B-C). 96.3% of caught prey were eventually eaten (52/54). Directionally appropriate mouth pointing, but not folding, is blocked if the subumbrella is cut between the mouth and the margin, suggesting directional communication between the two structures (Figure 2C).

**Figure 2:**
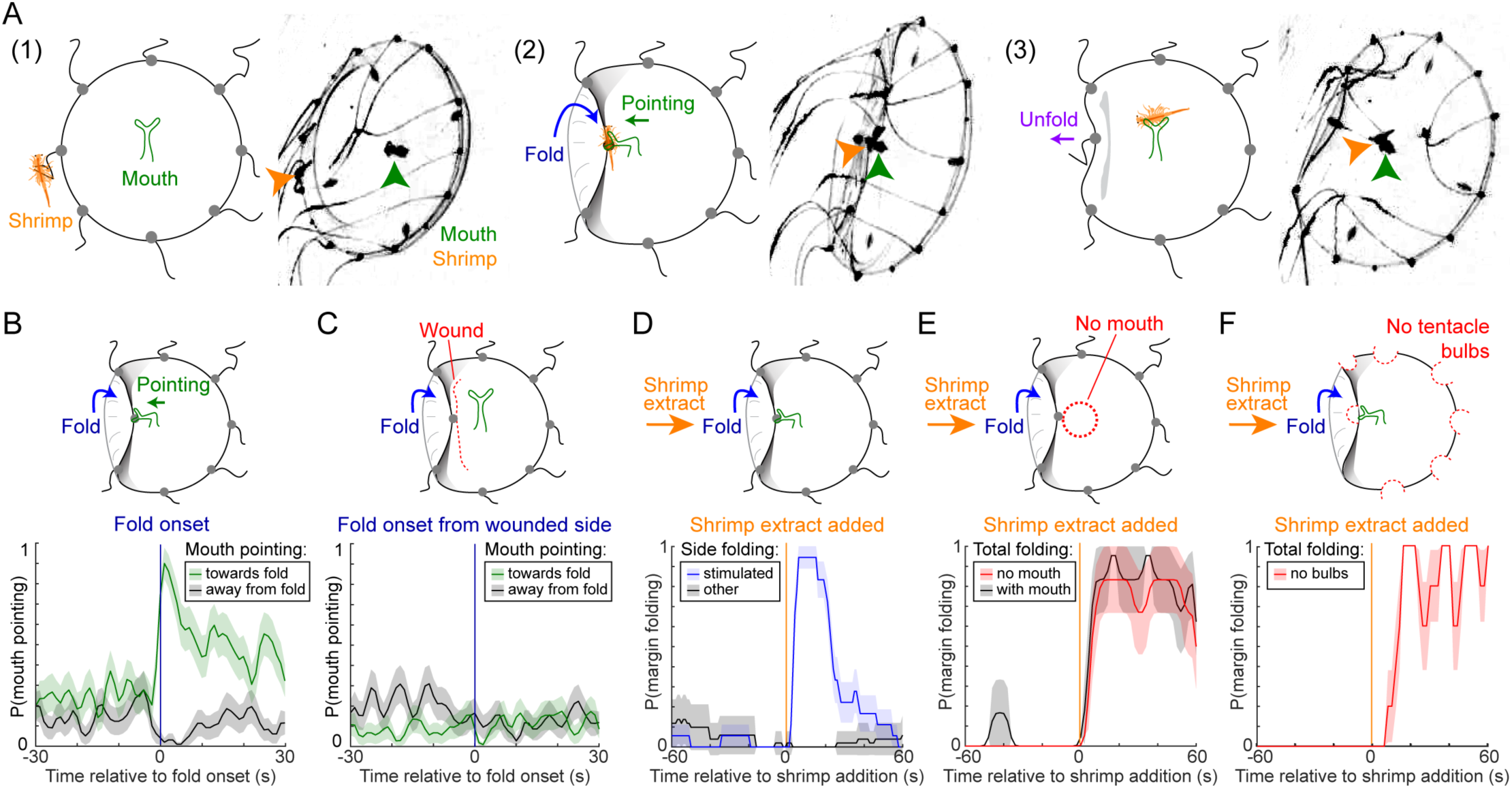
Margin folding is a coordinated, directionally targeted behavior. (A) Example video frames and illustrations showing the behavioral sequence of margin folding to pass prey from a tentacle to the mouth. First, a brine shrimp is captured by a tentacle (‘1’, left panels). Next, the corresponding portion of the margin folds inwards towards the mouth, and the mouth points in the direction of folding (‘2’, middle panels). Lastly, the prey is transferred to the mouth, and the margin relaxes back to the open position (‘3’, right panels). (B) Timing of mouth pointing towards the inward-folding margin (green line) but not in the opposite direction (grey line). These data, obtained from the unwounded side of the animals shown in (C), also serve as an internal control. Note that the mouth still points in the wounded direction spontaneously (baseline before fold onset). In all plots in this figure, data are mean ± SEM. n=52 folding events from 5 animals. (C) Mouth pointing requires communication with the margin. The subumbrella was wounded on one side of the animal. Mouth pointing towards the folding margin on the wounded side is eliminated (green line; cf. B). (D) Margin folding is triggered by local application of shrimp extract (blue line). Gray line represents a different quadrant. n=18 trials from 2 animals. (E) Margin folding does not require the mouth. Stimulus-triggered averages showing extract-triggered margin folding with (grey line) or without (red line) a mouth (n=6 each). (F) Margin folding does not require tentacle bulbs. Stimulus-triggered average showing margin folding by animals from which the tentacle bulbs have been removed. (n=5). See also Figure S1.

We observed that *Clytia* performed margin folding spontaneously without visible food present or stimuli provided (13.7±15% of time observed, mean±standard deviation; Figure S1D; Supplemental Video 2). This “margin folding behavior” is therefore used both for passing large prey items captured in the tentacles, and to bring the margin to the mouth in other contexts, possibly for cleaning. During margin folding epochs in which no prey were present in the tentacles, the mouth could often be seen interacting directly with the folded-in margin at regions located between the tentacles (Figure S1D, left; Supplemental Video 2).

Chemical but not mechanical stimuli were sufficient to trigger margin folding behavior (Figure S1E). Chemically induced margin folding was directionally appropriate when chemosensory cues (brine shrimp extract) were delivered locally to a spatially restricted segment of the margin (Figure 2D; Supplemental Video 3).

We next tested the necessity and sufficiency of different body parts for margin folding using surgical manipulations. Although communication between the margin and mouth during margin folding is necessary for mouth pointing (Figure 2B-C), surprisingly, we found that following surgical excision of the mouth, the body swims, captures prey in its tentacles, and tries to pass prey items to the hole where the mouth formerly was (Supplemental Video 4). The mouth-less umbrella also performed margin folding in response to chemosensory stimuli (Figure 2E), which could also trigger folding in the absence of tentacles and tentacle bulbs (Figure 2F).

Removal of other body parts revealed a similar theme: excised tentacles captured food and retracted, and the excised mouths continued to perform a number of behaviors autonomously, including directional pointing, prey capture, and eating (Figure S1F; Supplemental Video 5). Wedge-shaped strips of *Clytia* containing a tentacle at one end and the mouth at the other were able to perform the complete food passing behavior autonomously (Figure S1G; Supplemental Video 6), indicating that this behavior does not require an intact umbrella. Consistent with previous studies (Romanes, 1885; Passano, 1973; Quiroga Artigas et al., 2018; Takeda et al., 2018), further experiments demonstrated that this autonomy is a more general organizational principle that applies beyond food passing behavior. For example, small pieces of the margin continue to swim, and isolated gonads will spawn following the onset of light (Supplemental Video 5). Together, these experiments confirm a modular functional organization of *Clytia* medusae, in which autonomously functioning body parts, each containing their own behavioral control systems, interact in a coordinated manner to achieve higher-level, organismal behaviors.

These observations raise the question of the neural mechanisms that coordinate different body parts in a radially symmetric organism to achieve directionally oriented behaviors. As a first step towards addressing this question, we chose to focus on how asymmetric, vectorial folding of the bell is implemented in the umbrellar module. The neural elements associated with this module include the circumferential nerve rings at the margin and the subumbrellar nerve net; the mouth and tentacles are not required (Figure 2E-F).

### Identification and genetic targeting of RFamide-expressing neurons as a candidate population controlling margin folding behavior

Margin folding is expected to utilize contraction of the subumbrellar radial muscle, which is overlaid by the subumbrellar nerve net, which contains approximately 100-200 neurons in a 5mM animal (Figure 3A-B; Figure 1D-H; Satterlie, 1985). Using scRNAseq, we previously identified 14 subpopulations of neurons in the entire jellyfish, most of which express a distinct neuropeptide (Chari et al., 2021). Among these, arginine-phenylalanine-amide (RFamide)-expressing neurons have previously been identified in the subumbrellar nerve net in *Clytia* (Mackie et al., 1985), and were suggested to be involved in radial muscle contraction, including a proposed role in mouth pointing during feeding, in other jellyfish species (Grimmelikhuijzen, 1983; Grimmelikhuijzen and Spencer, 1984; Mackie, 2003; Satterlie, 2008, 2002; Spencer, 1989; Weber, 1989).

**Figure 3:**
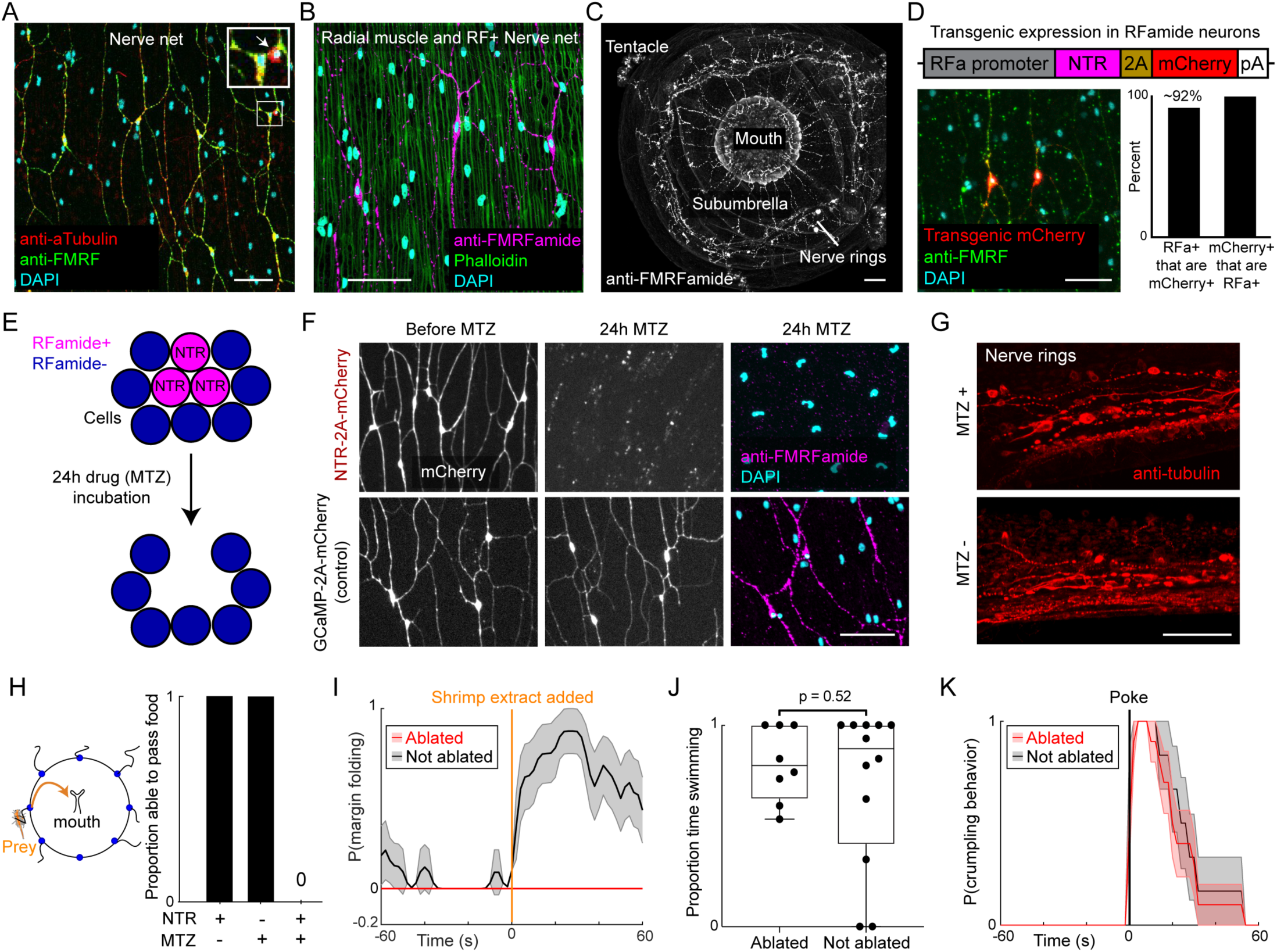
RFamide neurons are necessary for margin folding. (A) The majority (>80%) of nerve net neurons (red) are FMRFamide immunoreactive (green, “RFamide neurons”). Inset shows higher magnification of the white boxed area, with an RFamide-negative neuron indicated by a white arrow. Percent overlap calculated from n=3 locations, each from 4 animals. (B) RFamide neurons (magenta) are oriented along, and appose, the radial muscle (green). (C) Whole jellyfish showing distribution of RFamide neurons. Note radial orientation of the nerve net. (D) Transgenic cassette (top) showing upstream genomic fragment from *Clytia* RFamide precursor gene (“promoter”) (grey) driving Nitroreductase (NTR) and mCherry expression. Image illustrating mCherry overlap with anti-FMRFamide staining and quantification are shown below. n ≥ 3 locations from each of 4 animals. (E) Schematic illustrating scheme for conditional genetic ablation of RFamide neurons in transgenic jellyfish. MTZ, metronidazole. See text for details. Adapted from Curado et al., 2008. (F) Effect of MTZ incubation on RFamide neurons in NTR-expressing (top) and control non-NTR-expressing (bottom; see Fig. 4) transgenic animals. mCherry labeling (left and center) demonstrates that loss of FMRFamide antibody staining (right) reflects cell ablation, not loss of peptide expression. Scale ∼50µm. (G) Intact neurons in the nerve rings of RFamide::NTR-2A-mCherry animals following incubation in MTZ, demonstrating specificity of ablation. Scale bar 50µm. (H) Proportion of animals that successfully passed food to the mouth within 5 minutes of prey capture. NTR transgenic with MTZ addition, n=10 animals. Controls, n=6 each. (I) Stimulus-triggered average comparing margin folding between experimental (NTR transgenic + MTZ, red line) and control (NTR transgenic without MTZ, grey line) animals after shrimp extract was added to the water. n=8 each. Data are mean ± SEM. (J) RFamide neurons are not necessary for swimming. Plots show proportion of time swimming in NTR transgenics with (MTZ+, n=12) or without (MTZ-, n=8) MTZ treatment, 30-seconds/each. (K) Stimulus-triggered average of crumpling behavior following a poke delivered to the subumbrella. NTR transgenics with (red line, n=10) or without (grey line, n=6) MTZ treatment; mean ± SEM. Scale: 50µm. Further information in Figure S2.

Consistent with these earlier studies, we found by immunostaining that RFamide neurons make up the majority of nerve net neurons (∼80%, Figure 3A-B). These subumbrellar RFamide neurons are radially oriented with varicosities apposed to radial muscle fibers (Figure 3B; Figure S2A-D). In contrast, RFamide-negative neurons in the nerve net are generally smaller and lack clear radial orientation (Figure 3A, arrow; Figure S2A-D). RFamide-expressing neurons are also located in the tentacles, nerve rings, and mouth (Figure 3C, Figure S2E-G) but, unlike in the nerve net, they are not the dominant population overall (estimated ∼10% of total neurons in *Clytia* by scRNA-seq; Chari et al., 2021).

In order to investigate the function of the RFamide system, we established transgenesis in *Clytia* and used it to genetically ablate RFamide neurons. Transgenesis had not previously been established in any cnidarian with a free-swimming medusa (jellyfish) stage. Briefly, we microinjected plasmid and Tol2 transposase into eggs (Figure S2H; Koga et al., 1996; Ni et al., 2016), established stable F1 transgenic lines, and maintained them as clonal polyp colonies (see Methods). Colonies can be maintained indefinitely and expanded by vegetative propagation, and release clonal transgenic medusa (Figure 1C; Figure S2H). In order to target and ablate the RFamide neurons, we cloned the 6.6kb of genomic DNA immediately upstream of the open reading frame of the *Clytia* RFamide precursor gene and used it to regulate expression of a construct containing nitroreductase (NTR; Bridgewater et al., 1995; Tabor et al., 2014; White and Mumm, 2013) and mCherry (Figure 3D). This genomic fragment successfully drove strong and specific mCherry expression in the subumbrellar nerve net (∼92% of RFa^+^ neurons targeted, 100% of targeted neurons were RFa^+^, Figure 3D), affording genetic access to this population.

### RFamide neurons are required for margin folding

We next asked whether RFamide neurons are necessary for margin folding behavior, using the enhanced potency NTR system for cell ablation (Curado et al., 2008; Tabor et al., 2014). In this system, the NTR protein is genetically targeted to a cell type of interest; addition of the drug Metronidazole (MTZ) causes autonomous ablation of that cell type, as NTR converts MTZ to its toxic form intracellularly (Figure 3E). This system has been widely used and is highly efficient, cell type-specific, and inducible (Pisharath et al., 2007; White and Mumm, 2013).

24-hr incubation in MTZ successfully caused ablation of RFamide neurons in the RFamide::NTR-2A-mCherry transgenic line (Figure 3D), as was clear both from the loss of transgenic mCherry+ neurons (Figure 3F, left and center panels) and from loss of RFamide antibody staining (Figure 3F, right panels). Other neuronal populations were intact, demonstrating that ablation of neurons was not widespread (Figure 3G). No ablation was observed in controls with either MTZ or NTR components omitted (Figure 3F, left column and bottom row).

Remarkably, RFamide neuron ablation caused a complete loss of the ability to pass food when brine shrimp were applied to, and captured by, the tentacles of experimental animals (Figure 3H). RFamide neuron ablation also prevented margin folding behavior in response to chemosensory stimulation with shrimp extract (Figure 3I), which can occur in the absence of tentacles (Figure 2F). Thus, for both folding to pass prey caught in the tentacles, and for chemically induced folding, there was a complete loss of inward flexing of the margin.

We next examined other behaviors to determine whether the effect of RFamide neuron ablation was specific to margin folding. We quantified swimming behavior, which stops during margin folding (Figure S1A), and defensive “crumpling”, which uses the radial muscle to uniformly pull the margin inwards in response to a noxious stimulus (Hyman, 1940). Both swimming and defensive crumpling remained intact following neuronal ablation (Figure 3J-K). These experiments demonstrate that RFamide neurons are necessary for margin folding in multiple contexts, but are dispensable for other prominent umbrellar behaviors.

### RFamide neurons are active during margin folding behavior in multiple contexts

We next sought to determine how the activity of RFamide neurons relates to *Clytia* feeding and other behaviors, by imaging subumbrellar RFamide neurons expressing the calcium indicator GCaMP6s and mCherry (RFamide::GCaMP6s-2A-mCherry; Figure 4A; Chen et al., 2013). This transgenic line had a similarly high efficiency and specificity of targeting RFamide neurons as the NTR transgenic line (∼94% of RFa^+^ neurons targeted, 100% of targeted neurons were RFa^+^). *Clytia* have at least four endogenous green fluorescent protein (GFP) genes, expressed at different times and locations (Fourrage et al., 2014). In order to improve GCaMP imaging of the nerve net, we used CRISPR/Cas9 knocked out GFP1, which is highly expressed in the subumbrella (Momose et al., 2018) in the RFamide::GCaMP-2A-mCherry genetic background (Figure S3A).

**Figure 4:**
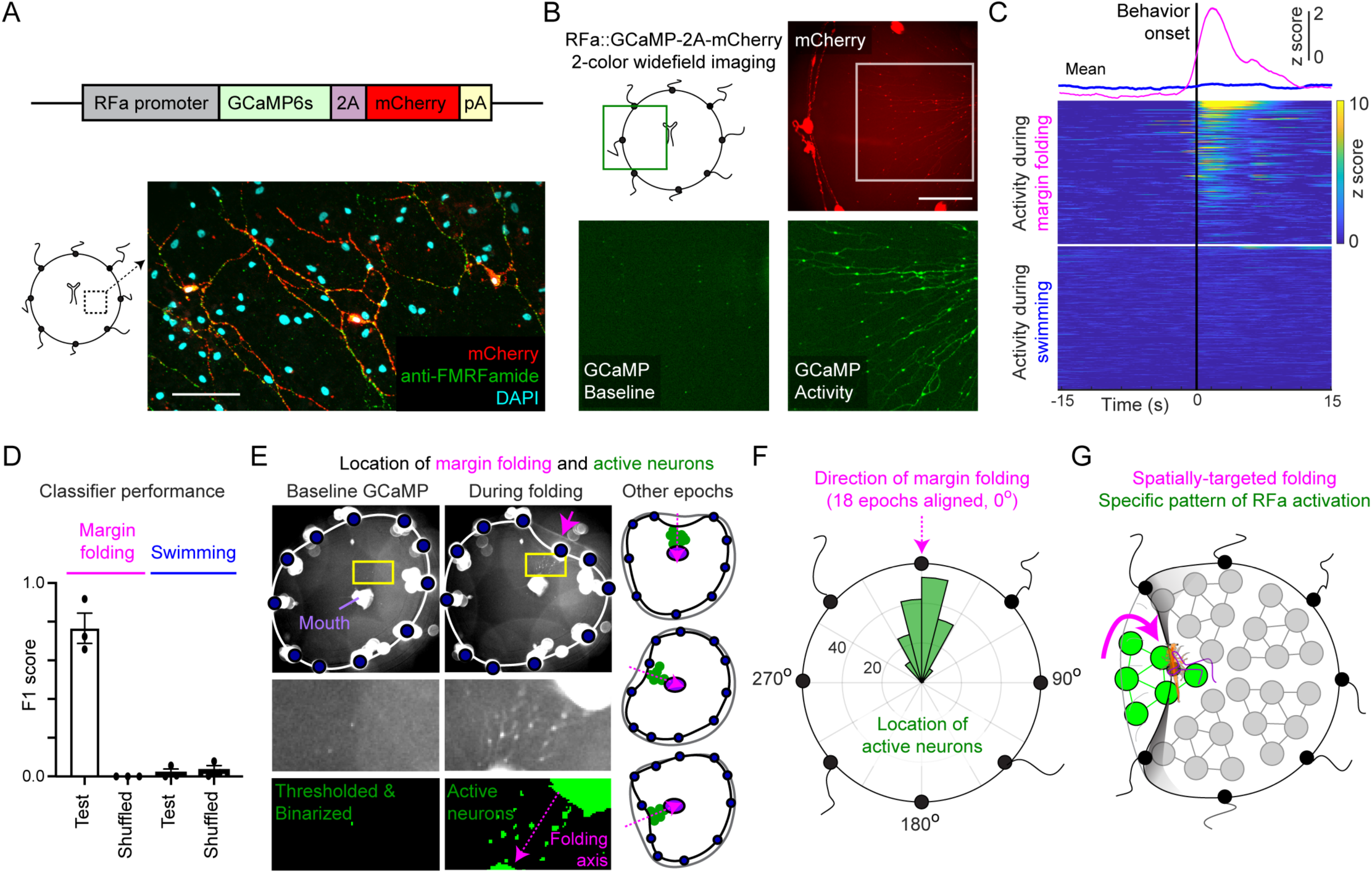
Activity of RFamide neurons is spatio-temporally correlated with margin folding. (A) Transgene structure (top) and expression (bottom) of GCaMP6s and mCherry in RFamide neurons. ∼94% of RFamide neurons targeted, 100% of targeted neurons were RFamide+. Scale: 50µm. (B) Example video frames from 2-color, widefield imaging showing mCherry (upper right) and GCaMP signal (bottom panels); magnified view of inset box in top panel. Scale: ∼1mm. (C) Behavior-triggered average of GCaMP activity during margin folding behavior (magenta) versus swimming (blue). Top traces show mean activity and heatmaps show the individual units. n=222 neuron-event pairs (margin folding) and 371 neuron-event pairs (swimming) from 4 animals. (D) Performance of 3-way Random Forest classifiers in predicting margin folding or swimming behaviors using population neural activity. Dots represent individual animals. (E) RFamide neurons are active during naturalistic behavior at the location of folding. Examples of activity during baseline (left column) and a folding event (right column) in minimally restrained animals. Bottom 4 panels are magnified views of yellow boxed regions (top). Annotated example frames (right) indicate active neurons (green dots) during folding events from different directions (magenta arrow). Blue dots: tentacle bulbs, purple dot: mouth. Shrimp extract diffusely present in the water (non-directional). FOV is ∼3.5mm^2^. (F) Polar histogram showing the location of active neurons (green bars) relative to the location of inward folding (magenta arrow). Multiple folding events from different directions were aligned to 0°; inset units are number of neurons (n=175 neurons from 18 folding events). (G) Schematic illustrating spatial location of RFamide neuron activity during margin folding. A radially oriented ensemble of neurons is active at the epicenter of the fold. See also Figure S3

We performed wide-field, two-color, *in vivo* calcium imaging in RFamide::GCaMP6s-2A-mCherry transgenic jellyfish, using preparations with different degrees of animal restraint that allowed for different behavioral and neural signals to be extracted (Figure 4B; Figure S3B). In one preparation, animals were minimally restrained and could behave naturalistically, allowing us to relate responding neurons to behavior (Figure 4E-F; Supplemental video 7). In another, animals were highly restrained (by agarose-embedding), allowing for extraction and analysis of high-quality GCaMP traces from single neurons (Figure 5). In a subset of experiments, there was both sufficient restraint to extract GCaMP traces and sufficient freedom of movement to measure animal behaviors (Figure 4C-D). In all cases we used the mCherry channel (collected simultaneously) to help segment neurons and correct for any motion artifacts in the GCaMP signal (Figure S3F; S4E).

**Figure 5:**
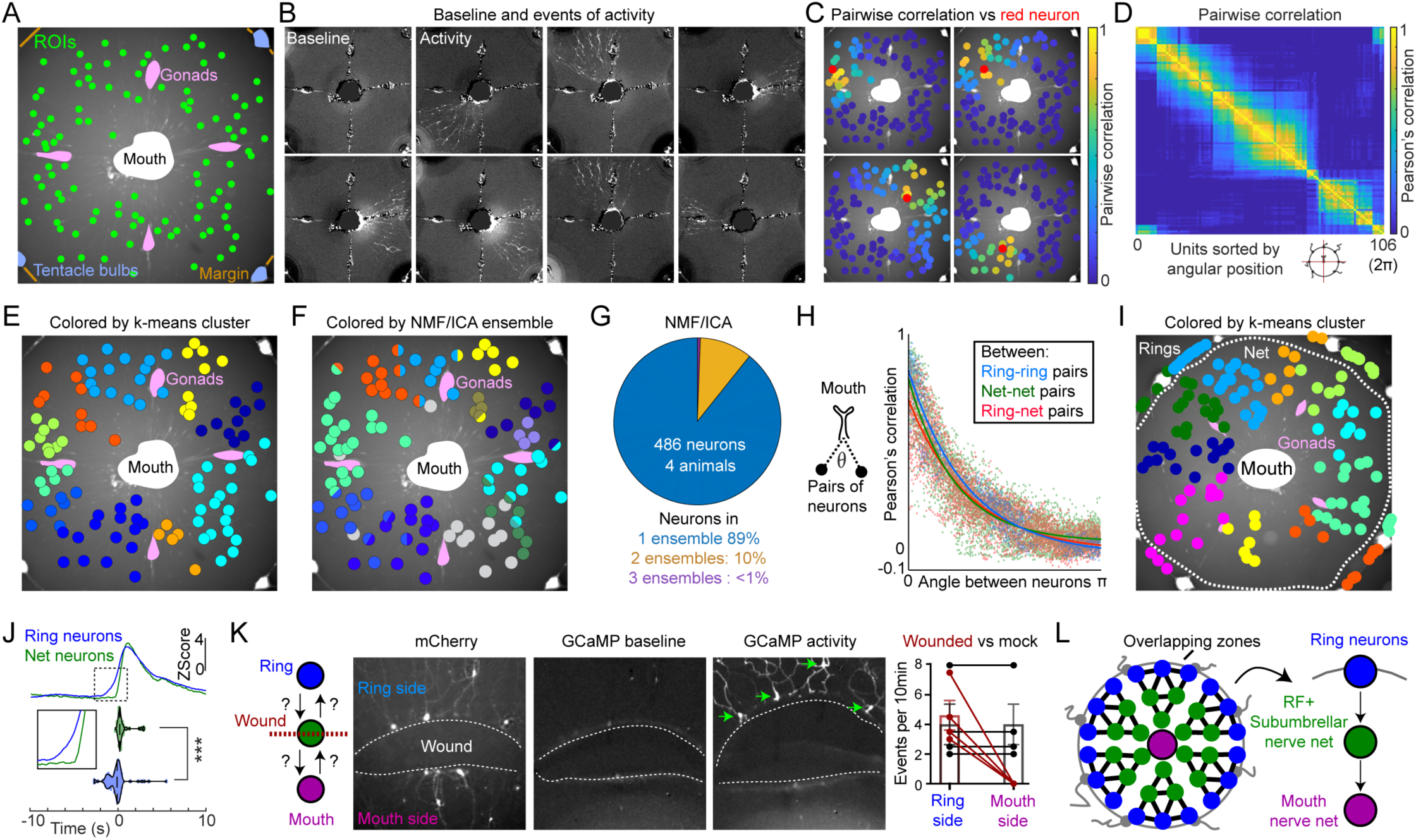
Functional parcellation of the umbrellar RFamide network. (A) Example field of view (FOV) for GCaMP imaging of spontaneous activity in highly restrained jellyfish. Center of ROIs (green circles) are superimposed on mCherry image. FOV is ∼3.5mm^2^. (B) Example frames of GCaMP imaging showing synchronous firing of neuronal ensembles. Baseline fluorescence (upper left) and representative spontaneous activity (remaining panels). (C) Each panel shows an example neuron (red) from the same animal with all other neurons colored by the pairwise Pearson’s correlation of their activity versus that example. (D) Pairwise Pearson’s correlation of neuronal activity (i.e. time of GCaMP peaks), sorted by angular position relative to the mouth. (E) k-means clustering of neurons based on their activity, with neurons colored by cluster membership. (F) NMF/ICA analysis, with neurons colored by ensemble membership. Multi-colored circles indicate membership in more than one ensemble. Neurons not assigned to ensembles are grey. (G) Percent of neurons that participate in 1, 2, or 3 ensembles using NMF/ICA. n=486 neurons from 4 animals. (H) Relationship between pairwise activity correlation and angle between neurons in the net or ring relative to the mouth. (Tau: Ring-ring: 0.97, Net-net: 0.75, Ring-Net: 0.99). (I) Ring neurons cluster with their corresponding net domain. FOV with neurons colored by k-means cluster membership. Same animal as analyzed in (H). (J) Nerve ring activity precedes nerve net activity. Top traces, mean ring activity (blue line) aligned to the onset of net activity (green line, t=0). Inset shows magnification of boxed region. Violin plots (below) show onset times relative to t=0 for individual neurons to cross a threshold z-score (1.5). Onset time difference is significant (p < 0.0001, data from n=4 animals). (K) Wounding prevents spread of activity from margin to mouth. A wound was generated in the subumbrella prior to imaging spontaneous activity (left diagram). Example images show mCherry (left) and GCaMP (right) channels. Green arrows indicate active neurons. Quantification (far right) shows GCaMP events per 10min on either the ring or mouth side of wounded (red) or control (black) animals (n=4 each). (L) Diagram showing model in which ensemble activity is organized into partially overlapping, radially oriented, wedge-shaped zones that tile around the animal (left). Within ensembles, information flows unidirectionally from the rings through the net towards the mouth (right). See also Figures S4 and S5.

We first chose experiments from the subset in which both behavior and GCaMP could be measured, where both margin folding and swimming were displayed in single imaging sessions, and compared behavioral epochs to neural activity (Figure 4C-D; Figure S3C). Consistent with our NTR ablation experiments, we found that RFamide neurons are strongly activated at the onset of margin folding but not during swimming (Figure 4C; Figure S3D). We trained a 3-way classifier to predict quiescence, swimming, and margin folding using RFamide population neural activity, and found that margin folding could be identified with high accuracy while swimming behavior could not be distinguished from quiescence (Figure 4D; Figure S3D). Principle components analysis (PCA) indicated that the largest fraction of variance was dominated by activity during margin folding (Figure S3E). These data provide strong evidence that RFamide neurons are involved specifically in margin folding behavior.

In order to examine the pattern of activation broadly across the nerve net under more naturalistic conditions, and to validate the findings above, we performed GCaMP imaging in a minimally restrained preparation while they performed margin folding in three contexts: (1) spontaneously, i.e., no stimuli given (Figure S1D), (2) with shrimp extract uniformly present in the water, or (3) with brine shrimp fed to the tentacles. Due to the high degree of movement in this setting, we were unable to extract clear traces of RFamide neuron activity from the entire recording; instead, we manually extracted body shape and active neurons during folding events. We found that RFamide neurons have a specific pattern of activation during margin folding that is similar in both the spontaneous and the two evoked contexts: at the onset of margin folding, there was activation of a spatially restricted, radially oriented wedge of nerve net neurons located at the epicenter of the folding event (Figure 4E-F; Figure S3F-G; Supplemental Video 7). These findings demonstrate that RFamide neurons are specifically correlated with margin folding in multiple, naturalistic contexts, and reveal a stereotyped pattern of neural activation corresponding to the time and position of each folding event (Figure 4G).

### Functional subdivisions within the umbrellar RFamide network

Our experiments in minimally restrained animals indicated a clear pattern of RFamide neuronal activity during margin folding, with a radially oriented ensemble of neurons activated at the epicenter of a folding event, aligned to the direction of folding (Figure 4F-G). This is particularly notable because our *a priori* expectation was for a diffuse ‘nerve net’ organization, in which the spread of activity through the network would be isotropic and unstructured. To confirm this structure, we performed whole organism imaging of the subumbrellar RFamide system in animals that were restrained in agarose (Figure 5A). Our preliminary observations indicated that spontaneous activation of the nerve net occurs under these conditions.

We found that large ensembles of RFamide neurons exhibit synchronous, spontaneous activity. These ensembles appear as radially oriented, wedge-shaped populations that stretch between the margin and the mouth (Figure 5B-C; Figure S4A-E; Supplemental Video 8). This was consistent with the pattern observed in minimally restrained animals performing both spontaneous and stimulus-induced behavior (Figure 4). This radial organization is evident in the pairwise correlations between neural activity over time when neurons are sorted by the angle between them relative to the mouth (Figure 5C-D, S4F).

When clustered based on their activity using k-means, RFamide neurons form striking spatial groupings that appear to roughly tile the animal (Figure 5E). Neighboring clusters of neurons were more likely than distant ones to activate simultaneously (Figure S4G). Because k-means forces neurons into single clusters, to capture variation in the ensemble membership of individual neurons across events, we additionally used Non-negative Matrix Factorization (NMF) followed by Independent Components Analysis (ICA) to identify ensembles (Figure 5F-G; Hyvärinen and Oja, 1997; Lopes-dos-Santos et al., 2013; Marčenko and Pastur, 1967; See et al., 2018). Remarkably, of neurons assigned to ensembles, the vast majority participated in only a single ensemble (∼89%), with far fewer participating in 2 (∼10%) or 3 ensembles (<1%; n=489 neurons from 4 animals; Figure 5G). Overlapping neurons were primarily observed at ensemble boundaries (Figure 5F). No structure was evident when using traces extracted from the mCherry channel as a control for motion or imaging artifacts (Figure S4H).

Interestingly, we found that activity in the nerve ring showed a spatial organization coordinated with that of the nerve net. Nerve ring neurons were highly correlated, and clustered, with their corresponding local, radial region of the nerve net, rather than more broadly within the rings, and were uncorrelated with both distant ring and net neurons (Figure 5H-I; Figure S4I-L). Together, these analyses reveal spatially segregated domains of correlated activity within the umbrellar RFamide system, in which partially overlapping yet largely distinct populations of neurons are arrayed around the animal. These radially organized activity domains are composed of both ring and net neurons, suggesting that domain affiliation, rather than membership in the ring or net, is the primary organizational principle of the umbrellar RFamide system (Figure 5L).

### The rings act upstream of the net as information flows from the rings towards the mouth

We next investigated the directionality of information flow between the RFamide neurons in the nerve rings and net. Our previous finding that the mouth is not necessary for margin folding (Figure 2E) led us to hypothesize that margin folding is initiated by activity in the nerve rings. Supporting this, we found that excising the margin (containing the rings, Figure 1D) leads to a loss of folding behavior in response to chemosensory cues (Figure S5A-B).

We next analyzed data from GCaMP imaging experiments in which both the ring and net were in the field of view, to determine whether a direction of activity propagation could be inferred from neural activity. Strikingly, we observed that nerve net activation always had a nerve ring correlate, but that nerve ring activity did not always coincide with an event in the nerve net (Figure S5C). Further, we found that the nerve ring is active before the nerve net during recordings of spontaneous activity (Figure 5J). Together, these observations are consistent with the idea that the rings may act upstream of the net to initiate margin folding.

We tested the model of inward information flow directly by generating wounds in the subumbrella, midway between the margin and the mouth, and monitoring whether nerve net activity could be observed on the ring side, the mouth side, or both, using GCaMP imaging (Figure 5K). Neural activity on the ring-side of subumbrellar wounds was common, but we never observed net activation on the mouth-side of wounds (Figure 5K, Supplemental Video 9), consistent with the relative timing of GCaMP activity between ring and net (Figure 5J). Thus, under these conditions, information flows through the subumbrellar RFamide nerve net from the rings inward towards the mouth (Figure 5L).

### Compartmentalized activity in the RFamide system reflects selection, not induction

Our previous analyses suggest a functional organization of the *Clytia* nerve net in which RFamide neurons form overlapping zones that are tiled around the animal (Figure 5L). This model largely arises from our finding that core ensembles of neurons repeatedly fired together (Figure 5). Such activity could be explained either by a *de novo* induction of ensembles at the site of prey capture (or other stimulus), or by a selection of the nearest ensemble from a pre-existing set that parcellate the net into stable domains. We therefore sought to determine whether these spatial zones are fixed or flexible (Figure 6A).

**Figure 6:**
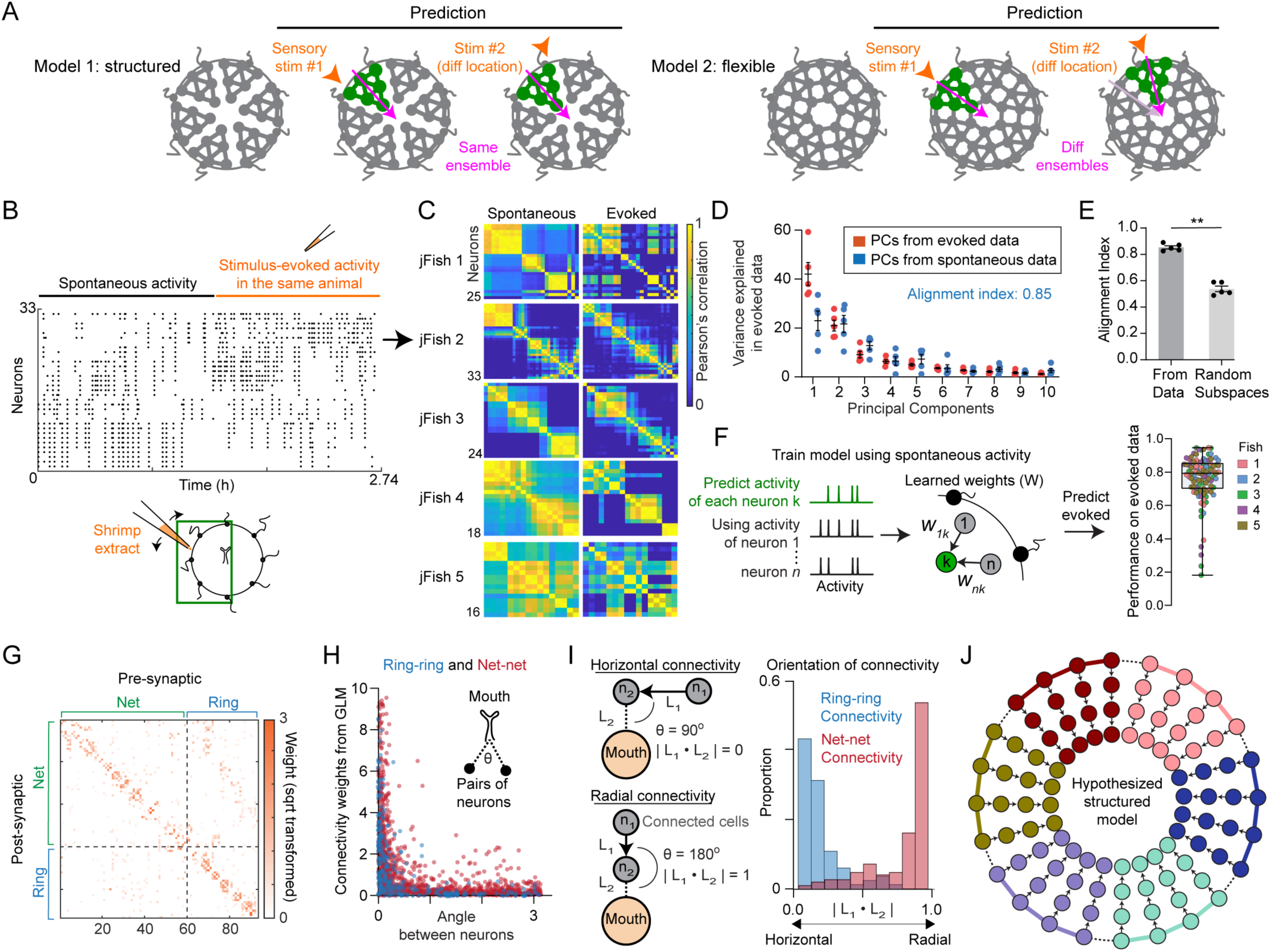
Ensemble activity arises from underlying structure. (A) Alternative models of system organization. Model #1 (left, “structured”): activity selects from a pre-existing set of ensembles. Model #2 (right, “flexible”): a new ensemble is generated at each stimulus location. (B) Example data showing spontaneous followed by evoked activity in the same session. Dots indicate time of a GCaMP peak. Diagram shows experiment and approximate FOV. (C) Pairwise correlation during spontaneous (left panels) and the matching evoked (right panels) epochs. Example data in panel B corresponds to jellyfish ^#^2. (D) Variance explained in evoked data using its own principle components (PCs, orange) or those from spontaneous activity (blue). The first 5 PCs from spontaneous activity explain 85% of the variance in evoked activity. Dots here and in panel E represent each of the 5 animals in (C). (E) Alignment indices, using the first 5 PCs from spontaneous activity to explain variance in evoked activity. Index is significantly higher than using PCs from random subspaces sampled in a data-correlation aligned manner (p = 0.0079, Mann-Whitney Test, see Methods). (F) Generalized linear models (GLMs) were trained to predict the activity of each neuron given the activity of every other neuron in the same jellyfish, using only the spontaneous activity epoch. Evoked neural activity could be predicted with high accuracy using these spontaneous activity-based GLMs (right, R^2^). (G) Example matrix of weights from GLMs trained on spontaneous activity. (H) GLM weights are highest for pairs of neurons with a small angle between them relative to the mouth. (I) The orientation of connectivity relative to the mouth (i.e. absolute value of the dot product) between pairs of neurons with strong connectivity (GLM weights >2) in the ring (“horizontal” connectivity) versus net (“vertical” connectivity). There is high vertical (radial) connectivity in the net, and horizontal (circumferential) connectivity in the rings (p = 3.5e-70, Kolmogorov–Smirnov test). (J) Working model of the organization of the umbrellar RFamide system, in which sparsely and radially connected nerve net neurons act downstream of horizontally connected nerve ring neurons, forming structures that tile the animal. See also Figure S6.

To distinguish these models, we imaged spontaneous activity in the nerve net followed by multiple rounds of chemosensory stimulation along the margin in the same animal and compared the structure in ensemble activity (Figure 6B-C; Figure S6A-B). If spontaneous nerve net activity reflects a constraint imposed by underlying structure, then spontaneous and evoked activity should be similar. If instead the system is more flexible, then new ensembles should be generated at the site of stimulus presentation.

We observed high subspace alignment between spontaneous and evoked epochs in all animals examined (85 ± 1.21%, mean ± SEM, Figure 6D-E), which was significantly different from randomly generated subspaces (p=0.0079). The subspace alignment metric (Elsayed et al., 2016; Yoo and Hayden, 2020) quantifies similarity as the alignment between the largest principal components of the spontaneous and evoked activity epochs (and vice versa). High subspace alignment indicates that the largest principal components of one epoch explain a large proportion of variance in the other, whereas low subspace alignment indicates distinct correlation structures. The high alignment observed here suggests that evoked activity does indeed select from an underlying stable distribution of intrinsic ensembles, supporting a model in which the RFamide system is organized into a fixed set of functional domains, possibly constrained by underlying connectivity.

As an independent method for comparing spontaneous to evoked activity patterns, we next trained generalized linear models (GLMs) to predict the activity of each neuron using the weighted activity of all other neurons (Figure 6F; Efron et al., 2004; Mishchenko et al., 2011; Pillow et al., 2008). A significant fitted weight indicates that a given “presynaptic” neuron has significant predictive power for the activity of a target “postsynaptic” cell. We found that GLMs trained on spontaneous activity and tested on held-out (spontaneous) data could recapitulate activity of a given postsynaptic cell with high accuracy (83 ± 1.3% mean ±SEM). More importantly, GLMs trained on spontaneous activity could also predict patterns of stimulus-evoked neural activity with similarly high accuracy (80 ± 1.4% mean ± SEM, Figure 6F), suggesting that they share a common constraint on their structure.

Lastly, we examined matrices of fitted GLM weights between RFamide neurons, which can be taken as a proxy for functional connectivity (Mishchenko et al., 2011). Interestingly, we observed that our models used only sparse, local weights between neurons (Figure S6C). To more confidently estimate weights, we additionally trained GLMs on the long, spontaneous recordings described in Figure 5 (Figure 6G-I; Figure S6D). In all animals examined, GLMs were able to recapitulate neural activity in held out test data with high accuracy (76 ± 1.2% mean ± SEM, Figure S6D). As above, our models generated sparse, local connectivity between neurons (Figure 6G-H).

Ensemble activity has both a radial and a circumferential dimension – i.e., inward flow from the rings towards the mouth, and the width of the ensemble, respectively (Figure 5L; Figure S4E). We therefore examined the relationship between weights among connected neurons and the angle of those connections relative to the mouth. Interestingly, within-net connectivity was primarily oriented radially whereas within-ring connectivity was primarily horizontal (circumferential), relative to the radial axis (Figure 6I; p = 3.5e-70). This modeling suggests that radial and circumferential components of ensemble structure may reflect the intrinsic connectivity schemes of the net and rings, respectively.

Taken together, the data and analyses presented here support a model in which the RFamide subumbrellar network and associated ring have a deterministic spatial structure underlying ensemble activity, comprised of partially overlapping, radial domains that tile the umbrella. This parcellation emerges from local interactions between sparsely and radially connected “strands” of RFamide neurons in the nerve net that act downstream of neighboring cells in the circumferentially connected nerve rings (Figure 6J). Local stimulation of the ring by captured prey selects the nearest radial domain for activation; this in turn facilitates oriented folding and transfer of food from the margin to the mouth.

## Discussion

In this study, we introduce *Clytia hemisph*aerica as a new model for systems neuroscience and perform the first systems-level interrogation of the nervous system of any jellyfish species using genetic tools. Like other medusozoans, *Clytia* have a distributed, modular nervous system organization in which body parts are able to perform behavioral subfunctions autonomously, but are coordinated in the intact organism to produce higher-level behaviors. As a first step towards understanding the neural basis of such coordination, we investigated the neural control of margin folding behavior, which is used to pass food from the tentacles to the mouth. Using genetically targeted ablations and calcium imaging in transgenic jellyfish, we reveal that such oriented, asymmetric behavior reflects an unexpected functional parcellation of the umbrellar nervous system.

### A new model organism for systems neuroscience

New techniques for sequencing and gene editing have dramatically lowered the barriers to entry for establishing new model systems, presenting the opportunity to match organisms to specific questions and to explore the diversity of life (Jourjine and Hoekstra, 2021; Laurent, 2020). In neuroscience, these advances have intersected with new, genetically encoded optical techniques for measuring and manipulating neural activity, together with advances in microscopy, offering the opportunity to functionally image the entire nervous system in small, transparent animals (Ahrens and Engert, 2015). With the exception of *Hydra* (a hydrozoan polyp; Dupre and Yuste, 2017; Szymanski and Yuste, 2019; Badhiwala et al., 2020; Wang et al., 2020), the only transparent model organisms currently used in systems neuroscience are *C. elegans* and zebrafish (*D. rerio*) larvae, both of which are bilaterians, although others are emerging (Bezares-Calderón et al., 2018; Chartier et al., 2018). Broadening the study of neural systems across phylogeny has utility that ranges from biomimetic engineering to revealing the possibilities, constraints, and principles through which nervous systems have evolved.

Here we have applied these tools to establish the first genetic jellyfish model for systems neuroscience, motivated both by its value from a comparative perspective and by the experimental tractability that it affords. Within jellyfish, we chose *Clytia hemisphaerica* because it is small, quasi-planar, and optically transparent, exhibits coordinated behaviors, and has a life cycle that facilitates aquaculture, transgenesis, and genetic crosses. Importantly, genetically modified strains can be asexually propagated indefinitely at the polyp stage, and easily distributed. Our work builds on pioneering efforts by Houliston and colleagues, who initially established *Clytia* as a model for studies of early development and evolution (Houliston et al., 2010). In this inaugural study, we have extended this toolkit by developing methods to generate stable, F1 transgenic jellyfish lines, and have used a cell type-specific promoter to investigate the role of a specific neuronal subpopulation in feeding behavior, via calcium imaging and genetic ablation. With these tools in hand, the potential uses of *Clytia* as a jellyfish model are wide ranging, from identifying conserved and derived features of nervous system organization and function to studies of neural development and regeneration (Sinigaglia et al., 2020).

### New insights into the systems-level neural control of jellyfish behavior

A misconception of jellyfish in popular culture is that they are relatively unstructured, gelatinous masses that do little more in the way of behavior than to propel themselves through the water via rhythmic, isotropic contractions of their bell (umbrella). Their nervous system has been correspondingly viewed as distributed and relatively unstructured, and therefore well-suited to mediate such simple, symmetric organismal movements. However, recent work is revealing previously underappreciated complexity in jellyfish behavior (Fossette et al., 2015; Kaartvedt et al., 2015; Lewis and Long, 2005; Nath et al., 2017). Here we show, using *in toto* calcium imaging of the umbrella in intact, behaving animals, that the *Clytia* RFamide system is functionally organized into a series of wedge-shaped, partially overlapping sub-networks defined by local connectivity rules. We provide evidence that this structure is employed to achieve local contractions of the umbrellar musculature, that are in turn used to vectorially transfer prey from the margin (which folds inwards at the site where food has been captured by the tentacles) to the central mouth. We suggest that this organization is critical to allow the organism to perform oriented, asymmetric behaviors and to coordinate interactions between otherwise autonomously functioning body parts. This insight could not have been achieved without population imaging of neural activity and subsequent computational analysis.

It is important to note caveats with our imaging approach, however. First, our high-resolution imaging analyses relied largely on data from infrequent, spontaneous displays of neural activity in highly restrained animals. We interpret this activity as occurrences of the activity pattern that normally underlies spontaneous margin folding behavior in unrestrained animals. In support of this interpretation, spontaneous activity in the nerve net without accompanying margin folding behavior was not observed in loosely restrained animals. Further, activity evoked by food in restrained animals had a similar pattern as spontaneous activity, and in restrained animals, flexing of the radial muscle could be observed accompanying spontaneous activity bouts. Importantly, such spontaneous activity was not induced by restraint itself, since it could also be observed in free-swimming medusae. Whether such “spontaneous” activity is truly endogenously generated, or rather is triggered by microscopic particles or non-randomly distributed chemical cues, remains to be determined. While we cannot exclude that activity in highly restrained animals may exhibit subtle differences from the patterns in freely moving jellyfish, analysis of loosely restrained animals revealed a qualitatively similar regionalization of activation. Improvements in methods for fast volumetric imaging and image registration may eventually allow routine imaging of neural activity in intact, freely behaving *Clytia* (Kim et al., 2017).

Our modeling suggests that the observed structure in the *activity* of the network may emerge from an underlying structure in the *connectivity* of the system. This connectivity appears to be sparse and local, with radially oriented connections in the net and circumferential connections in the rings. Our findings reveal that the “overlapping zone” architecture in the subumbrellar neural net extends to neurons in the nerve rings, which we have suggested act functionally upstream of the net in food passing. In this view, activated domains in the ring would in turn activate neighboring wedges of radially connected nerve net neurons, leading to local contraction that brings the margin in contact with the mouth. Whether vectorial bending of the mouth towards the folding margin is also controlled by such subumbrellar wedges remains unknown, and will be an interesting question for future studies.

Although we have emphasized the zonal organization of the subumbrellar network, our data indicate that some neurons are shared between ensembles and exhibit flexibility in their ensemble membership. Indeed, this flexibility accounts for roughly 15% of both the overlap between ensembles and the variance when comparing spontaneous to evoked activity. While this variance may reflect technical and/or biological noise, it may also be an optimal solution to the problem of linking the precise spatial location of stimuli on the margin to activation of the corresponding muscles needed to transfer food to the mouth. Further studies may reveal additional flexibility in ensemble structure, depending on critical features of the stimulus, the behavioral context, or life stage.

### Role of the RFamide system in margin folding

We have used expression of the RFamide neuropeptide as a genetic marker to label a subpopulation of neurons and examine their role in a behavior at the systems level. RFamide-expressing neurons comprise over 80% of the neurons in the subumbrellar net. Our data indicate that the activity of these neurons is correlated with, and necessary for, margin folding behavior, with activity localized at the epicenter of the folding event. These results provide strong evidence that RFamide neurons are important for margin folding. However they do not exclude a role for non-RFamide neurons in this behavior.

Our data leave open the question of the precise role of RFamide neurons in the circuits that control margin folding. On one end of a spectrum, the RFamide system could operate as a self-contained, molecularly defined functional module to initiate and execute margin folding: nerve ring neurons could include both sensory and command neurons; nerve net neurons could be both motor neurons and relays to the mouth. On the other end of the spectrum, the RFamide population could serve an essential function, perhaps as interneurons, but not play an instructive role in action selection or execution of the behavior.

Moving forward, it will also be important to address the role of the neuropeptide itself. RFamide peptides in *Hydra* can act through ionotropic receptors and appear to be used in neuromuscular transmission (Assmann et al., 2014; Golubovic et al., 2007; Gründer and Assmann, 2015). This would be consistent with the idea that nerve net neurons have a motor neuron function, and that the RFamide peptide itself may serve to trigger contraction of the radial muscle. However, the peptide could also be used in a modulatory manner, perhaps acting in concert with the co-release of a classical transmitter. Future studies performing knockout and overexpression of the RFamide peptide and its receptors, imaging neuropeptide release (Ding et al., 2019), and other strategies, will be needed to address these questions.

Although we have focused in this study on the RFamide neurons in the umbrella (i.e. rings and net), RFamide neurons are also concentrated in the mouth and tentacles. An attractive hypothesis is that these neurons may be important for the functions of those additional body parts in food passing or other feeding behaviors. In that case, communication between parts could be achieved by virtue of co-expression of the RFamide peptide and its receptor at each stage in the relay. Alternatively, the RFamide population may play distinct and divergent roles in each body part. Future experiments imaging and manipulating neuronal populations in the tentacles and mouth will be needed to address these questions. Such studies may shed light on the larger issue of the mapping relationship between molecular cell types and behaviors, both across regions within an animal, and across different species.

### Behavioral and nervous system diversity and evolution in Hydrozoans

There is an extensive literature on feeding behavior in various jellyfish species. In hydrozoans, descriptions of food passing behaviors date back over a hundred years (Romanes, 1885; Hyman, 1940; Horridge, 1955; Miglietta et al., 2000; Mackie and Singla, 1975). The most detailed previous study of food passing behavior and associated neural mechanisms examined the hydrozoan *Aglantha* (Mackie, 2003). Interestingly, *Aglantha* uses pointing of an elongated mouth (“mouth pointing”), but not margin folding, to retrieve prey from the tentacles. Mouth pointing could be evoked by chemosensory stimulation of the margin independently of the tentacles, similar to our findings in *Clytia*. Interestingly, this behavior was found to be controlled by several discrete bundles of FMRFamide-containing axons that run between the margin and mouth (rather than the “nerve net” organization in *Clytia*). The presence of RFamide in the relevant nerve bundles may suggest a conserved role for the peptide in feeding behavior, independent of the details of its physical implementation.

Feeding behavior incorporating mouth pointing and margin folding, as described here in *Clytia*, has been described in related jellyfish species, such as the leptomedusa *Aequorea* (Satterlie, 1985) and multiple species of narcomedusae (Larson et al., 1989). However, other species of hydrozoans have been described as passing food to the mouth directly via tentacle bending, or by using simultaneous contraction of all of the radial muscle rather than by vectorial folding (Miglietta et al., 2000; Tachibana et al., 2020). These data suggest a rich diversity in the implementation of food passing behavior amongst jellyfish species. The evolutionary substrate for this diversity, and whether it involves modifications to the RFamide system, remain to be determined. As hydrozoan medusae are often small and transparent, with a range of interesting behavioral and morphological diversity beyond feeding, they may provide a fruitful platform for comparative evolutionary systems neuroscience moving forward (Koizumi et al., 2015; Satterlie and Spencer, 1983).

An additional opportunity for comparative neuroscience would be between *Clytia* and *Hydra,* which is also emerging as a systems neuroscience model (Dupre and Yuste, 2017; Szymanski and Yuste, 2019; Badhiwala et al., 2020; Wang et al., 2020; Bosch et al., 2017)*. Hydra* most closely resemble the polyp stage of *Clytia,* but have undergone a number of remarkable modifications, including loss of the polyp colony and movement into fresh water (Chapman et al., 2010; Leclère et al., 2019; Steele et al., 2011). These lifestyle changes have been accompanied by new behaviors in the *Hydra* lineage, such as movement by “somersaulting” (Han et al., 2018; Trembley, 1744). With the new technical capabilities for *Clytia* described here, comparison of the nervous systems of *Clytia* polyps and *Hydra* could provide insights into the evolution of such qualitatively new behaviors in the *Hydra* lineage.

Direct comparisons between *Hydra* (polyps) and *Clytia* medusae are more challenging, but interesting similarities and differences do emerge. Similar to our findings in the *Clytia* RFamide system, synchronous firing of distinct ensembles of neurons has been described in *Hydra*, with both spontaneous and stimulus-evoked activity (Dupre and Yuste, 2017; Badhiwala et al., 2020; Tzouanas et al., 2020). Differences appear as well, including the bidirectional propagation described in *Hydra* (Dupre and Yuste, 2017), which is unlike the unidirectional flow in the nerve net described here, though bidirectional flow may exist in other *Clytia* systems. Regardless of the particulars of this preliminary comparison, the continued development of multiple, complementary cnidarian neuroscience models has important implications for understanding the origin of nervous systems and principles of their evolution and function (Bosch et al., 2017).

### Decentralization and modularity of neural systems in organismal behavior

One of the most striking features of *Clytia* is its extreme functional modularity, with small pieces of the animal able to perform certain behaviors in isolation. This initially led us to posit that *Clytia* behavior may simply result from the physical coupling of autonomous modules; i.e. that *Clytia* are a sum of their parts. However, our early and outwardly simple finding that the mouth points in the direction of margin folding indicates at least some degree of coordination between parts during food passing. Further, our experiments indicate that wedge-shaped strips of *Clytia* containing a tentacle at one end and mouth at the other are able to perform the complete passing behavior, following prey capture by the tentacle. This suggests a hierarchical organization of functional modules, with super-modules that combine individual body parts able to carry out a coordinated behavior. In that case, organism-level behavior could emerge by duplicating and arraying these super-modules around the animal, to allow food passing to occur from any site on the margin. Whether new organism-level behaviors appear once such super-modules are combined is currently unknown. This nested organization, in which increasingly complex features emerge via hierarchical combinations of simpler functional modules, may be an important substrate for the evolution of complex behaviors, and a more general feature observed in complex systems.

In organisms that have undergone cephalization, modularity in the form of “brain regions” is widely observed across neural systems, likely providing important computational benefits. Nevertheless, most such organisms retain a peripheral nervous system. These peripheral systems exhibit a certain amount of functional autonomy, such as the enteric nervous system that controls contractions of the gut (Kandel, 2013; Luo, 2015). They also have regenerative capacity, a likely feature of ancestral nervous systems (Tanaka and Ferretti, 2009). *Clytia* have remarkable abilities to regenerate and recover behaviorally following injury (Kamran et al., 2017; Sinigaglia et al., 2020), and also are continuously integrating newborn neurons into their nervous system without disrupting organismal function (Chari et al., 2021). The local network interactions and modular systems that we have described here may facilitate such growth and repair. *Clytia* presents an exciting, genetically tractable platform to understand the mechanisms underlying such dynamics, which lie at the interface of development, regeneration, and neural systems.

## Acknowledgements

We thank J. Malamy, E. Houliston, R. Copley, J. Costello, S. Colin, and members of L. Goentoro’s Laboratory, particularly T. Basinger, for assistance establishing *Clytia* at Caltech and performing initial experiments. We thank J. Malamy for first introducing B.W. and D.J.A to *Clytia,* X. Da and X. Wang for technical assistance, G. Mancuso for administrative assistance, C. Chiu for laboratory management, S. Ekker and C. Daby for assistance and reagents while establishing the Tol2 system, A. Collazo and the Caltech Biological Imaging Facility for imaging assistance, T. Chari, J. Gehring, and L. Pachter for scRNAseq analysis, and J. DeGiorgis who captured Supplemental Video 2. We thank E. Houliston, J. Parker, D. Prober, T. Karigo, G. Mountoufaris, Y. Ouadah, S. Stagkourakis, A. Vinograd, and B. Yang for feedback on the manuscript. This work was supported in part by the Caltech Center for Evolutionary Science, the Whitman Center of the Marine Biological Laboratory in Woods Hole, MA, a Howard Hughes Medical Institute Fellowship of the Life Sciences Research Foundation (to B.W.), the National Institute Of Neurological Disorders and Stroke of the National Institutes of Health under Award Number K99NS119749 (to B.W.), and by the National Institute Of Mental Health of the National Institutes of Health under Award Number K99MH117264 (to A.K.). The content is solely the responsibility of the authors and does not necessarily represent the official views of the National Institutes of Health. We thank the CRBM Villefranche (FR 3761) for marine facilities, biological resources, and a travel grant to B.W., which is supported by EMBRC-France, whose French state funds are managed by the ANR within the Investments of the Future program under reference ANR-10-INBS-02. T.M was supported by Agence Nationale de la Recherche (ANR), ANR-17-CE13-0016 (i-MMEJ). A. N. is supported by a National Science Scholarship from the Agency of Science, Technology and Research, Singapore. D.J.A. is an Investigator of the Howard Hughes Medical Institute.

## Author contributions

B.W and D.J.A. conceived of the project and wrote the manuscript, with input from T.M., A.N., A.K., and B.H.. B.W., D.J.A, and B.H. designed and performed histology, behavior, and imaging experiments, and B.W., D.J.A, and T.M. designed and performed experiments establishing transgenesis. B.W., A.N., and A.K. analyzed the data; A.N. performed NMF/ICA, subspace, and GLM analyses.

## Methods

### Animals

Culture of the *Clytia* life cycle was carried out in accordance with published protocols (Lechable et al., 2020), with the exception of the culture tank design. Our circulating system at Caltech uses the same overall flow design as in Lechable et al., 2020, but uses modifications of zebrafish tanks (Pentair) to provide either high flow for polyp slides held in glass slide racks (Fisher, cat#02-912-615) in small tanks, or low flow for jellyfish maintained in large tanks. Jellyfish tanks use a curved plastic insert with a large nylon mesh near the back overflow outlet to keep jellyfish in the tank; a constant speed 5rpm DC motor (Uxcell) attached to a dimmer switch (to tune the rotation speed) is then used to create a constant circular current. A slow drip from the circulating system into these tanks allows for water turnover without risking sweeping the jellyfish through the outlet. Jellyfish used for transgenesis, and all polyps, were maintained in these circulating systems, while smaller jellyfish were maintained in beakers. In beakers, current was generated using stirring with a DC motor, as above, attached to the lid of a multi-well tissue culture plate. All artificial sea water for culture and experiments was made using Red Sea Salts (Bulk Reef Supply, cat# 207077) diluted into building deionized water to 36ppt.

Unless otherwise indicated, behavior experiments were performed using sexually mature animals of the Z4B strain of *Clytia,* which are female. Transgenesis was performed by crossing Z4B females to Z13 males. For generating F1 lines, depending on whether the F0 was male or female, it was backcrossed to either Z4B or Z13.

### Histology

For antibody staining, *Clytia* were fixed for 2h at room temperature in 4% PFA in filtered artificial sea water (Red Sea Salts, Bulk Reef Supply, cat# 207077, diluted into building deionized water to 36ppt). They were then washed 3x in PBS followed by blocking for 1h in PBS with 0.1% Triton (PBST) with 10% normal donkey serum (NDS). Animals were then incubation for 1-3 nights in primary antibody with 5% NDS in PBST at 4-degrees. Primary antibodies used in this study were: anti-FMRF (Immunostar, cat#20091), anti-Tyrosine Tubulin (sigma T9028-100UL), and 647-conjugated anti-aTubulin, clone DM1a, (Millipore Sigma cat# 05-829-AF647). Following primary antibody incubation, one short (∼5min) and then 3-4 long (∼30min+) washes were performed in PBST, and then animals were transferred into secondary antibodies and/or Phalloidin-488 (Thermo Fisher, cat#A12379) for 2h at room temperature or overnight at 4-degrees. Secondary antibodies used in this study were donkey anti-rabbit conjugated to Alexa Fluor 647 or 488 (Jackson ImmunoResearch). Animals were stained with DAPI (BD cat#564907) and mounted onto glass slides for imaging. For staining shown in Figure 1E, animals were dehydrated stepwise into methanol and then rehydrated prior to the blocking step. Quantification of overlap in Figure 3D was from at least 3 separate locations/each from at least 3 animals. In situ hybridization was carried out as described in (Chari et al., 2021), including the RFamide probe, which was the same as the one used in that study. Imaging of histological specimens was carried out using an Olympus FV3000 confocal.

### Cloning

To generate the Actin::mCherry plasmid used to establish transgenesis, codon-optimized mCherry (Che-mCherry) cDNA was designed using COOL (Yu et al., 2017). The ACT2 promoter was cloned from upstream of a non-muscle actin gene (XLOC_011689) using primers: TTTGCTGCGTACAACAACAACGACC and TCGACTTATGTCCTGATAGTTCGGA. The 3’UTR used in all constructs was from a different actin gene (XLOC_021750) and was amplified using primers: CCAACAGATGTGGATCTCCAAACA and ACTGGAAGCCTGAGTTCCATCAAA. This was assembled into the pT2AL200R150G backbone (Urasaki et al., 2006; licensed under MTA - N° K2010-008).

Other *Clytia* transgenesis constructs were based on the miniTol2 backbone, a gift from Dr. Stephen Ekker (Addgene plasmid # 31829). To generate RFamide::NTR-2A-mCherry, the RFamide fragment was amplified from *Clytia* genomic DNA using the following primers, ATCCCCATCCGCCATCTTTG, GTTGTGTTCTTTCTTGATTTGATGG, and inserted into the miniTol2 backbone using In-Fusion Cloning (Takara). This miniTol2-RFamide backbone was then used to insert different effectors, always using In-Fusion Cloning, following digestion with Spe1. To generate RFamide::epNTR-2A-mCherry and RFamide::GCaMP6s-2A-mCherry: epNTR was amplified from the pCS2-epNTR plasmid, a gift from Dr. Harold Burgess (Addgene plasmid # 62213); both GCaMP6s and the 2A peptide used in this paper was derived from AAV-hSyn1-GCaMP6s-P2A-nls-dTomato, a gift from Jonathan Ting (Addgene plasmid # 51084).

### Transgenesis

In order to establish and optimize transgenesis, we first chose a DNA fragment upstream of a *Clytia* actin gene that we found to drive strong, widespread expression in planula and in polyp tentacles, enabling accurate estimates of efficiency during the critical early life stages following injection (Figure S2H). Using Actin-mCherry to test strategies, we established a protocol that now enables routine, efficient transgenesis, using microinjection of Tol2 transposase protein together with plasmid DNA into unfertilized eggs. Collection of eggs and sperm, and microinjection, was carried out as previously described (Momose et al., 2018). Briefly, *Clytia* medusa spawn ∼2 hours after the onset of light. In order to collect eggs and sperm, animals were transferred into either dishes (for the females) or 6-well plates (for the males) within the first hour of the lights coming on. After spawning, eggs were immediately collected and injected with a mixture of 6.25ng/ul Tol2 transposase protein and 10ng/ul plasmid DNA using a Femtojet (Eppendorf). Pulled glass capillaries were used for microinjection (WPI, TW100F-4). Rather than use the ‘inject’ function on the Femtojet, injections were carried out by puncturing eggs and then using the backpressure in the capillary to fill to the desired volume. Tol2 protein was produced by Creative Biomart using a plasmid generously provided by Dr. Stephen Ekker (Ni et al., 2016). Protein was then stored at -80 as 5ul aliquots and thawed directly prior to injection.

Immediately following injection – and within an hour of spawning – eggs were fertilized and allowed to develop overnight into planula larvae, at which point they were transferred into a 12-well plate of sea water containing penicillin/streptomycin. This prevents early metamorphosis of the planula into polyps, which appears to be triggered by bacterial cues. On the second day following injection, we checked the expression of plasmids in the planula to ensure that they were capable of driving sufficient expression. Importantly, at this stage expression is not dependent on integration. Planula were then induced to metamorphose into primary polyps onto glass slides (Ted Pella, cat #260439) using a synthetic GLWamide neuropeptide (produced by Genscript; Lechable et al., 2020).

Primary polyps were hand-fed mashed shrimp until they began to grow into a colony. Mashed shrimp were generated by drawing brine shrimp into a 10ml syringe with a blunt-tipped needle attached and then expelling them while pressing the end of the needle against the bottom of a dish or beaker. Once colonies had 3 or more polyps, they were screened for transgenic expression, with all but the most highly and broadly expressing polyp colonies removed. These colonies were then allowed to grow until they began releasing jellyfish. Jellyfish were then collected, raised to maturity, and crossed to wildtype in order to generate stable F1 colonies. As issues can arise, possibly due to inbreeding, F1s were seeded sparsely on slides, and were screened for several criteria: strong expression of the transgene, strong polyp colony growth and health, and the ability to release healthy jellyfish that were able to reach maturity. F1 colonies were maintained as single clonal colonies per slide, and once the best colony was identified, the rest were thrown away and that colony was expanded. This allows for the same genetic background to be used across experiments, as these colonies then release clonal experimental jellyfish as needed.

### Behavioral analysis and NTR ablations

Behavioral analysis was carried out on videos acquired through an Olympus stereoscope (SZX2) using Flea3 or Grasshopper USB3 cameras from FLIR, and the manufacturer’s acquisition software (FlyCapture). For all behavior experiments, videos were manually annotated using the BENTO analysis suite (https://github.com/annkennedy/bento).

Mouth pointing shown in Figure 2B-C was assayed by pinning animals to sylgard-coated plates (Dow Corning) using stainless steel minutien pins (Fine Science Tools). This prevented the margin from getting close to the mouth during a folding event, which would have the potential confound of causing directional mouth pointing by direct sensory stimulation of the mouth. The subumbrella was then wounded only on the right side of the animal to compare pointing in the intact (Figure 2B) versus the wounded (Figure 2C) direction. Having the internal negative control of the wounded direction ruled out the possibility that the mouth and margin were responding to a shared, directional sensory stimulus, or that the folding of the margin itself was a directional sensory stimulus for the mouth (e.g. by creating fluid flow). Videos were acquired at 15fps, and mouth pointing events 30s before and after a margin folding event were manually annotated using these videos. If the mouth was leaning in one direction at the start of the epoch, that lean was treated as a baseline for further pointing. All margin folding events were spontaneous (i.e. no stimuli were delivered) to avoid possible shared sensory stimulation of the margin and mouth.

Shrimp extract used in all stimulation experiments was generated by homogenizing brine shrimp using a syringe and a blunt tipped needle followed by filtration through a 40µm cell strainer. Experiments in this paper used either this 40µm filtered extract, or extract that was passed through an additional 0.2µm filter. Mechanical stimuli in Figure S1E was delivered by gently touching the tentacles and margin using a glass Pasteur pipette. For experiments in which shrimp extract was used to trigger behavior in freely moving animals, animals were transferred into 6-well plates and 20ul of 40µm filtered shrimp extract was added to the well.

For the directional folding experiments shown in Figure 2D, animals were pinned to sylgard plates, as described above for mouth pointing experiments. ∼5ul of 0.2µm filtered shrimp extract was then pipetted directly onto either the top, bottom, left, or right portion of the margin. Margin folding events from all quadrants, and the locations of sensory stimulation, were then manually annotated and compared. For physical ablation experiments, body parts were cut off using spring scissors (Fine Science Tools) or, to remove the mouth, by creating hole-punches using a blunt-tipped needle.

For NTR ablations, animals were incubated overnight in 10mM Metronidazole (Fisher, cat# ICN15571005; Curado et al., 2008). Immediately prior to behavior experiments, animals were washed several times in clean artificial sea water. For crumpling behavior, the subumbrella was gently poked with a glass pipette and crumpling duration was manually quantified.

### GCaMP imaging acquisition, processing, and data analysis

For highly restrained imaging experiments, *Clytia* medusa were embedded in agarose, as follows. 3-4%, Type VII-A, low-melting point agarose (Sigma A0701-25G) was first made in artificial sea water, with particular care to avoid evaporation. Agarose was then aliquoted and kept in a heat block set at 50-degrees until ready to use. Single tubes were then removed from the heat block and vortexed occasionally until reaching nearly room temperature. Medusa were then added into the tube, gently mixed, and then rapidly transferred to a glass-bottomed dish (Ted Pella, 14036-20). They were then quickly spread out to make them as planar as possible before being briefly touched to a cold object to rapidly cool and harden the agarose. Agarose was then covered with a thin layer of mineral oil (Sigma M5310-1L) to avoid evaporation during imaging experiments.

For minimally restrained imaging experiments, animals were placed into a glass depression slide, and a coverslip was gently placed on top, using small amounts of Vaseline applied to each corner of the coverslip as a spacer. The coverslip was then gently pressed down to create a small chamber in which the jellyfish could still perform behaviors. Experiments in which both behavior and GCaMP traces could be analyzed were from animals in which agarose embedding was attempted but they were poorly restrained. There was consistently a relatively small portion of the animal in the field of view in these experiments.

GCaMP imaging experiments always had synchronous acquisition of the red and green channels. This was achieved using an Olympus BX51WI microscope with two Photometrics Prime95B cameras connected using a W-View Gemini-2 Optical Splitter from Hamamatsu. Acquisition was controlled with Olympus Cellsens software. Images were acquired with downsampling during acquisition (to 600×600 pixels) and then further processed using ImageJ software (NIH). First, images were re-sized to 400×400 pixels. An average Z-projection was then performed, and each frame in the video stack was divided by this projection to generate a normalized intensity over time for each pixel. If needed, a spatial filter was applied using ImageJ’s Bandpass Filter function to remove spatial light artifacts, e.g., from movement of the mouth. Regions of Interest (ROIs) were then circled using the ImageJ ROI Manager, and the average pixel intensity with, and location of, these ROIs was then exported for further processing using Matlab (Mathworks). We found that the data was characterized by having large, discrete events of activity, and that by using events rather than raw traces we could more accurately represent the activity by removing any underlying noise in the trace. Events were detected in one of two ways, either using Matlab’s “findpeaks” function or using spike inference from the CNMF_E software package (Zhou et al., 2018) with parameters individually adjusted for each trace. Following spike detection, the first spike in a bout of inferred spikes was used as the timing of the event. In both cases, these events were then smoothed for several frames on either side to avoid false-negatives in correlated activity based on assignment of events to single neighboring frames. These smoothed events were then used for downstream analyses shown in Figure 5 and 6. For the behavior-triggered averages, classifiers, and other analyses shown in Figure 4, and the relationship between ring and net neurons shown in Figure 5, smoothed and z-scored GCaMP traces were used rather than extracting events, in order to more accurately retain timing information.

Classifier analysis was performed in Matlab (Mathworks). The first 1500 frames were used for training, with the remaining frames held out for testing; this ranged from 15%-36% of the data in the training set, depending on the length of the recording. 3-way, Random Forest classifiers had 60 bags and used out-of-bag prediction to distinguish between epochs of margin folding, swimming, and quiescence. SVM classifiers did pairwise classification between margin folding and quiescence, or swimming and quiescence. For the loosely-restrained experiments in Figure 4, and imaging-with-stimulation experiments shown in Figure S6, neurons that responded to the stimulus and body shape were manually annotated using the ImageJ ROI Manager, and Adobe Illustrator, by comparing the frame before initiation of a folding event to a frame during the folding event (Figure 4E; Figure S3F). The polar histogram shown in Figure 4 was generated using the XY coordinates of these annotated responders relative to the folding axis, determined by the location of the mouth and the center of the inward fold. For analysis of the timing of ring versus net activity, groups of net neurons and their corresponding ring neurons were extracted, and, for each event, the activity of ring neurons was aligned to the onset of a net event, which was found using the mean of that group of net neurons. To quantify the relative onset time within these “net-triggered averages”, any neuron that crossed a threshold of z-score > 1.5 within 5-seconds before the identified net onset, or 10 seconds after, was included, and that time to crossing was used for comparison. For the wounding experiments in Figure 5K, wounds were generated in the subumbrella using either a scalpel or forceps, and the number of events on each side of the wound was quantified. Mock wounds were unwounded regions in similar locations; a line was drawn on the video after acquisition using ImageJ, and the number of events on either side of the line was quantified.

For detection of cell ensembles using NMF/ICA, the method was based on a method described in (Lopes-dos-Santos et al., 2013) and used in (See et al., 2018). To determine the number of ensembles in each jellyfish, principal component analysis (PCA) was applied to data, processed as described above (extracted and smoothed events), to obtain the eigenvalue spectrum. Eigenvalues that exceed the upper bounds of the Marčenko-Pastur distribution (Marčenko and Pastur, 1967) are deemed significant and represent the number of detected ensembles. We determined this bound using a statistical threshold based on surrogate data. To obtain this threshold, we shuffled the time bins of each neuron independently in order to destroy their temporal relations while maintaining the distribution of events. The eigenvalues of correlation matrices obtain from shuffled data were used to construct a null distribution. Eigenvalues of the original data matrix that are larger than the 99th percentile of the distribution of maximal eigenvalues computed from shuffled data are regarded are significant (Lopes-dos-Santos et al., 2013). From the identified number of significant eigenvalues, we used non-negative matrix factorization (NMF) to optimize ensemble detection so that we maximize the number of overall neurons included in any ensemble. The resulting factors from NMF were then processed using the fast independent component analysis (fastICA) algorithm (Hyvärinen and Oja, 1997). The resulting independent components (ICs) represent the contribution of each neuron to each ensemble. We validated ensemble membership by applying the NMF/ICA method on shuffled data described above. This process was performed 100 times to obtain a normal distribution of IC weights for shuffled data. A neuron was considered a member of an ensemble if its IC weight was larger than 2 times the standard deviation of the IC weight distribution from shuffled data. With this cutoff, 79% of neurons were assigned to one or more ensembles.

For comparison of spontaneous and evoked neural activity spaces: to examine whether the structure of neural activity was similar during spontaneous and evoked epochs, we used an approach that compares the neural subspace occupied during the two epochs (Elsayed et al., 2016; Yoo and Hayden, 2020). We first performed PCA on the neural activity matrix obtained during each epoch. To examine the relationship between the two epochs, we projected activity during the evoked epoch onto principal components (PCs) obtained during the spontaneous epoch. We then quantified the percent of variance explained by spontaneous epoch PCs relative to the total variance of the evoked epoch (We also performed this analysis in reverse). We then defined the alignment index (*A*) as the ratio between the overall variance explained in an evoked epoch by its top 5 PCs (which explain >80% of variance) and the amount of variance of evoked data explained by spontaneous epoch PCs. This can be written as:

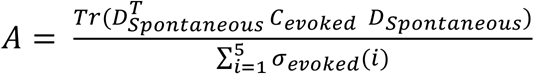

Where *D_spontaneous_* is the set of top 5 eigenvectors obtained by PCA in the spontaneous epoch. *C_evoked_* is the covariance matrix of the evoked epoch. The term *σ_evoked_*(*i*) is the i-th singular value of C_evoked. The term *Tr()* is the matrix trace which sums along the diagonal entry. Since the quantity in the denominator serves to normalize the alignment index, it ranges from 0 to 1, with 0 indicative perfect orthogonality between the subspaces and 1 indicating perfect alignment.

To obtain a statistical comparison, we generated random subspaces biased to the data covariance structure as described in (Elsayed et al., 2016). This was performed as a Monte-Carlo analysis applied to the covariance structure (*C*) from neural responses across all times (both spontaneous and evoked epochs). Random subspaces aligned to the structure of C are obtained as:

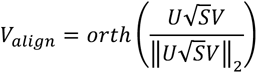

Where *U* and *S* are eigenvectors and eigenvalues of *C* respectively. *V* is a matrix with each element drawn independently from a normal distribution with mean zero and variance zero. *Orth(Z)* returns the orthonormal basis for a matrix *Z*. Thus, this procedure samples subspaces biased towards the space of neural activity, such that the sampled subspace will have the specified covariance matrix *C*. To calculate the alignment index of two sets of random subspaces 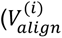 and 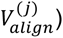 of dimension *d* =5, we calculate:

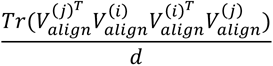

We repeat the random sampling procedure 1000 times to obtain a distribution of random alignment matrices for each jellyfish.

For the use of generalized linear models (GLMs) to infer functional connectivity between neurons in the spontaneous epoch (Mishchenko et al., 2011; Pillow et al., 2008), Gaussian-residual GLMs were fit for each neuron k using data from spontaneous epochs as:

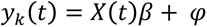

Where *y_i_* is the activity of target neuron *k* and *X* is the matrix of activity of all neurons in a given jellyfish except *k*. As a result, *β* is a vector of coefficient weights which can be interpreted as a proxy of the connectivity of all other neurons to neuron *k*. Lastly *φ* is an error term. We regularized our model with an additional non-negativity constraint. This was implemented using a non-negative least angle regression of least absolute shrinkage and selection operator (LASSO, Efron et al., 2004) to impose sparseness and non-negativity on model weights β, with the sparseness parameter determined using 10-fold cross validation.

To assess whether data from the evoked epoch could be predicted using model weights found from the spontaneous epoch, we evaluated model performance of GLMs for each cell using held-out test data from the evoked epoch. Model performance in Fig 6F is measured using the Pearson’s correlation coefficient (PCC) between predicted activity and ground truth evoked epoch data. GLMs were additionally trained on the recordings shown in Figure 5 to examine the connectivity matrix obtained from model weights. These recordings were longer and had more of the jellyfish in the field of view as compared to the more complex imaging followed by stimulation experiments. For this, we used the same GLM described above but assessed model performance on held-out test data from the same spontaneous epoch (rather than comparing to an evoked epoch), reporting model fit as the PCC between predicted activity and ground truth spontaneous epoch data. Model weights from GLMs of cells with a model fit R^2^ of at least 0.7 were used to construct the connectivity matrix in Figure 6G.

