## Supplemental Information for "Functional modules within a distributed neural network control feeding in a model medusa"

Supplemental Figure 1

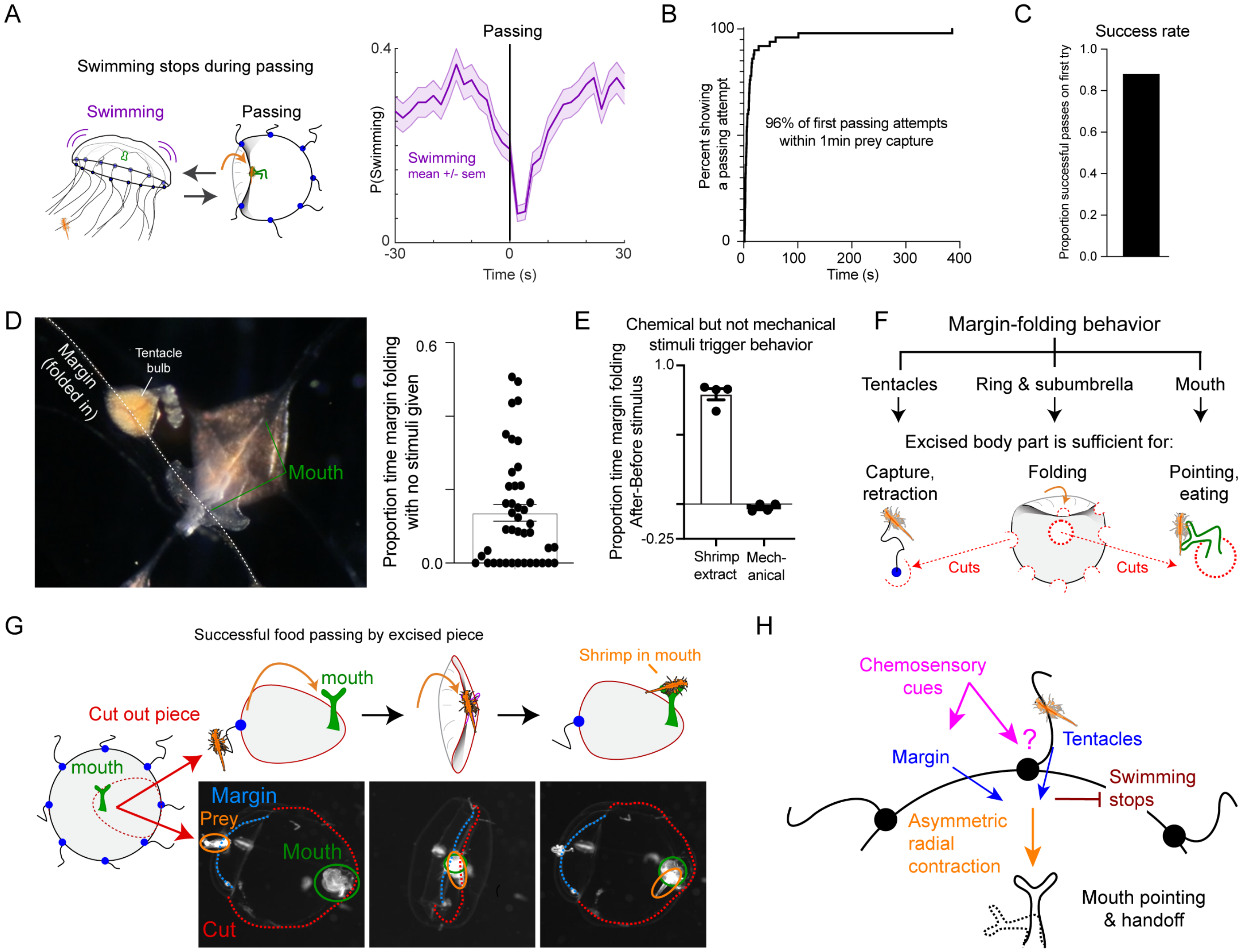

**Figure S1: Characterization of margin folding behavior**

Related to Figure 2.

(A) Swimming stops during margin folding. Mean ± SEM. n=218 passing events from 4 animals.

(B) Length of time between brine shrimp capture and attempted passing to the mouth. n=50 passing events from 17 animals.

(C) Success rate of passing prey from tentacles to the mouth during a margin folding event. n= 17 animals.

(D) Margin folding behavior can be observed with no visible prey present or stimuli provided. Panel on the left shows the mouth interacting directly with the folded-in margin (base and top of the feeding organ, “mouth”, indicated in green). Data on the right is pooled from 42 animals and shows the proportion of 2-minutes spent performing margin folding with no stimulus provided.

(E) The difference in time spent margin folding following stimulus presentation (2min after – 2min before). Chemosensory stimuli were mashed, filtered brine shrimp. Mechanical stimulus was given using a glass pipette applied to the tentacles and margin. Dots represent individual animals (n=4 each). Mean ± SEM is shown.

(F) Summary diagram showing that excised body parts are capable of performing subsets of the complete food passing behavior autonomously.

(G) Diagrams and example images showing an experiment in which a strip of the animal was cut out, with a tentacle at one end and the mouth at the other. When a brine shrimp was fed to the tentacle, this strip was able to perform the complete food passing behavior, handing the shrimp off to the mouth.

(H) Diagrammatic summary of margin folding behavior.

Supplemental Figure 2

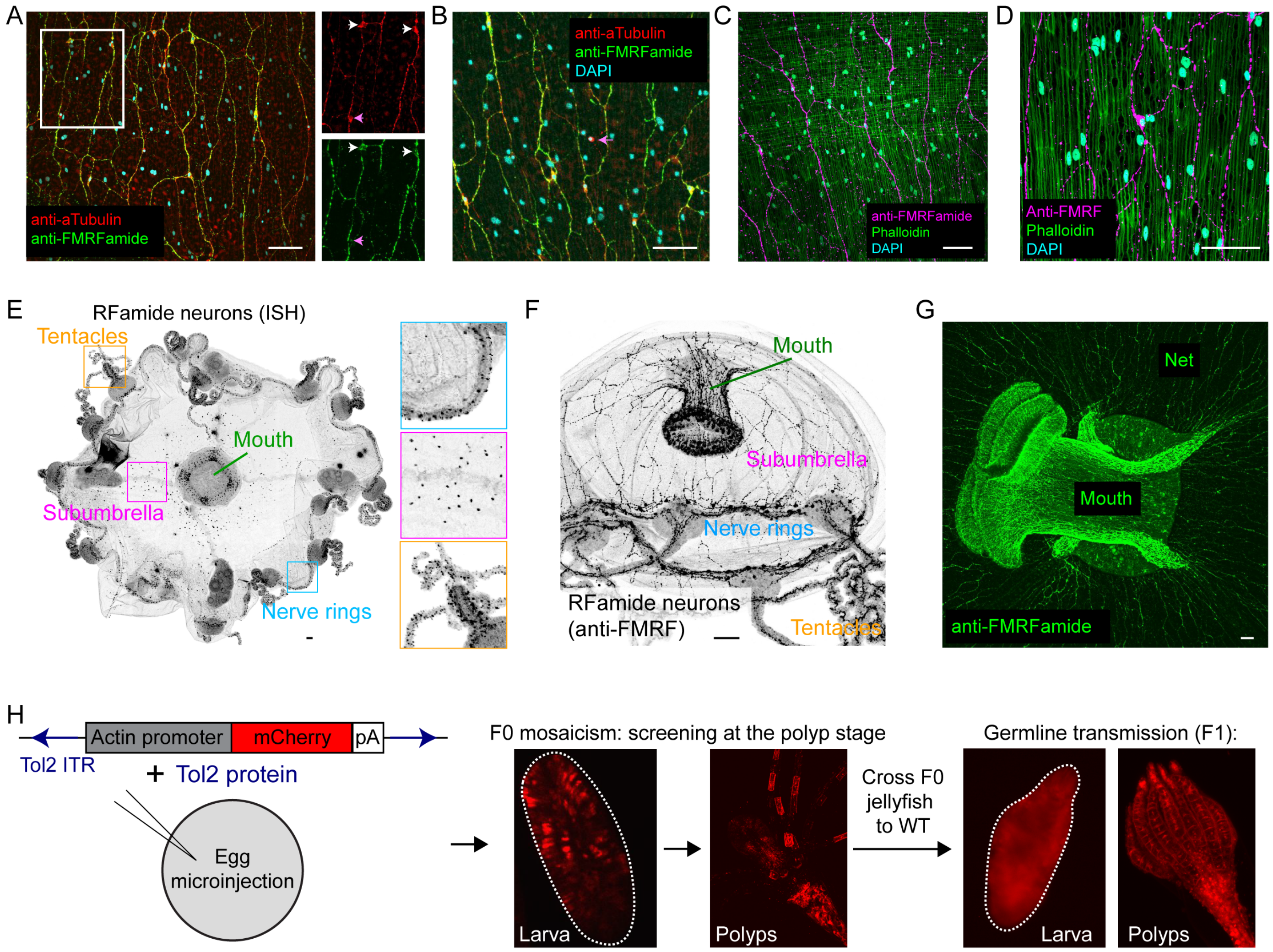

**Figure S2: Characterization of RFamide neurons and development of transgenesis**

Related to Figure 3.

(A-B) The majority (>80%) of nerve net neurons (red) are FMRFamide immunoreactive (green, "RFamide neurons"). White arrows indicate RFamide-positive nerve net neurons, and magenta arrows indicate RFamide-negative nerve net neurons. Small panels to the right of A are magnifications of the inset white box showing the separate color channels.

(C-D) RFamide neurons (magenta) are oriented along, and appose, the radial muscle (green). Circular, striated muscle is visible in the top half of the image in panel C, highlighting the radial orientation of the RFamide neurons and their association with radial muscle.

(E) In situ hybridization (ISH) for the RFamide precursor gene showing its distribution across the organism.

(F) The distribution of RFamide neurons across the organism (side view).

(G) The RFamide neurons of the mouth, and their association with the nerve net.

(H) Strategy for generating transgenic *Clytia*. Plasmid DNA containing Tol2 recognition sites was injected into eggs together with Tol2 protein (left). Larva were then induced to metamorphose into polyps, and polyp colonies were screened for the highest expression and lowest level of mosaicism (middle panels). Jellyfish were then collected from the best colonies and crossed to wildtype to generate stable F1 lines, which were re-screened and maintained as clonal colonies. See Methods for further details.

Scale: 50μm.

Supplemental Figure 3

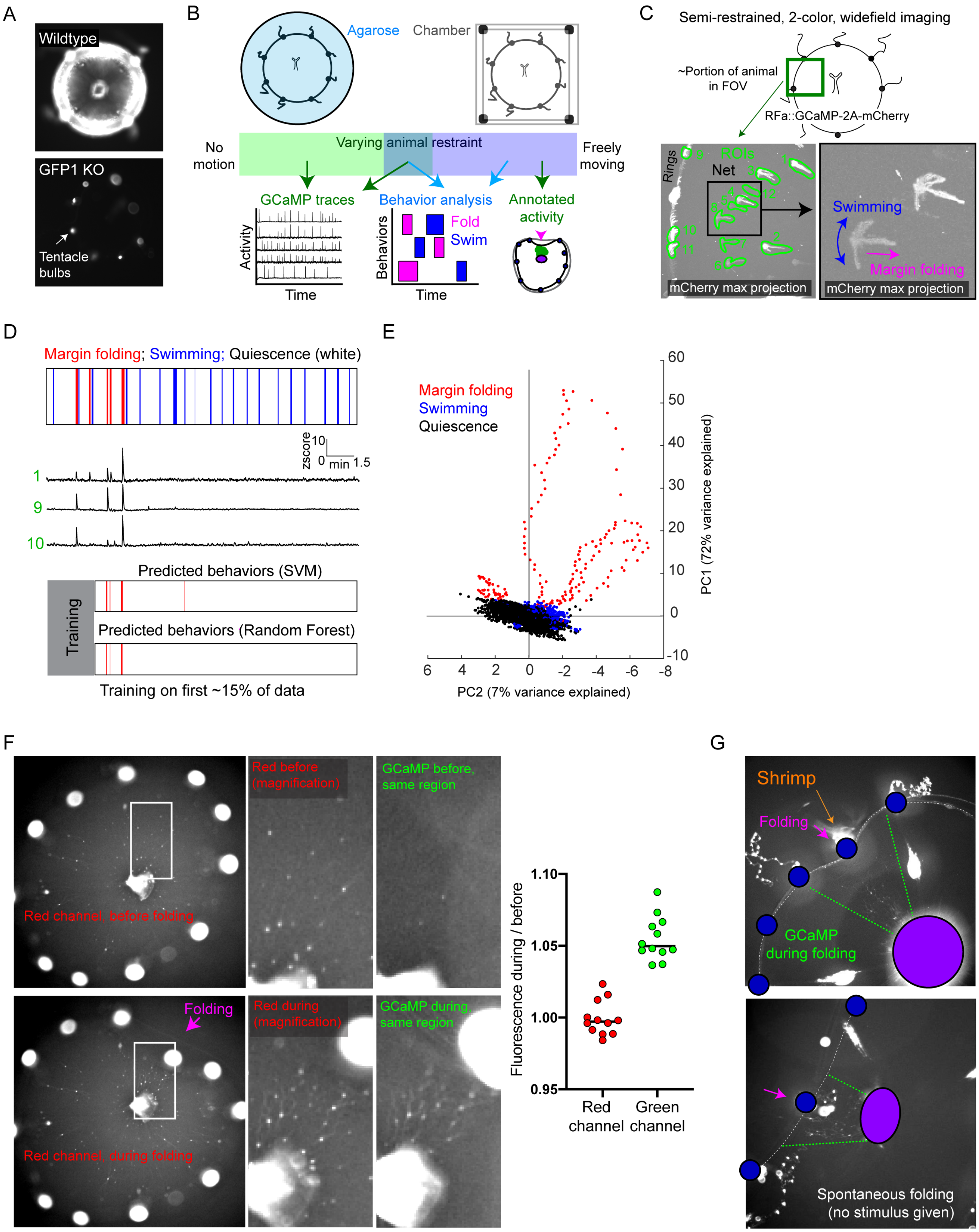

**Figure S3: RFamide neurons are specifically active during margin folding in multiple contexts**

Related to Figure 4.

(A) Images of wildtype (top) and GFP1 knockout (bottom) animals. Other GFP genes are expressed in the tentacle bulbs (arrow) and mouth.

(B) Overview of imaging experiments using varying degrees of animal restraint. On one end of the spectrum, highly restrained, agarose-embedded animals allowed extraction of high quality GCaMP traces. On the other, minimally restrained animals behaving naturalistically in small chambers allowed analysis of manually extracted GCaMP activity and body shape from individual frames. In a subset of experiments, we could both discern which behavior was being performed and extract GCaMP traces.

(C) Example of an experiment in which both behavior and GCaMP traces could be extracted. Top schematic shows approximate portion of the animal in the FOV. Bottom panels show the maximum projection of the mCherry channel with ROIs superimposed (left). The bottom right panel is a zoom-in of the black box in the left panel. Since this shows a maximum projection of mCherry, which is always acquired simultaneously during GCaMP imaging, the location of neurons over time is visible; side-to-side movement is indicative of swimming, while radial movement occurs during margin folding. FOV: ~3.5mm$^2$.

(D) Raster (top) shows annotated behaviors with example GCaMP traces below, where green numbers correspond to the ROIs in panel C. Bottom panels show classifier predictions using the first 15% of the data (greyed out) to predict the remaining behavior using population neural activity. Both classifiers were able to predict margin folding (red lines) but fail to predict swimming.

(E) The first two principle components over time of population neural activity, colored by the behavior being performed. Epochs of margin folding lead to trajectories, primarily out along the first principle component, while bouts of swimming and quiescence remain intermingled at the origin. Data corresponds to animal shown in C-D.

(F) Examples of GCaMP activity in a naturalistically behaving animal. Fluorescence in the simultaneously acquired red (left two columns) and GCaMP channels (right column) immediately before (top row) or during (bottom row) a margin folding event. GCaMP activation is visible in the green channel, but a similar change in fluorescence intensity is not observed in the red channel. Quantification shows fluorescence from frames during/before the folding event for ROIs manually circled around individual neurons in this field of view. Line represents the median. FOV here and in panel G is ~3.5mm$^2$.

(G) Example GCaMP frames during margin folding behavior showing that the characteristic pattern of RFamide neuron activation is also present during margin folding in other contexts, i.e. during passing of a shrimp captured in the tentacles (top) or during a spontaneous margin folding event (bottom).

Supplemental Figure 4

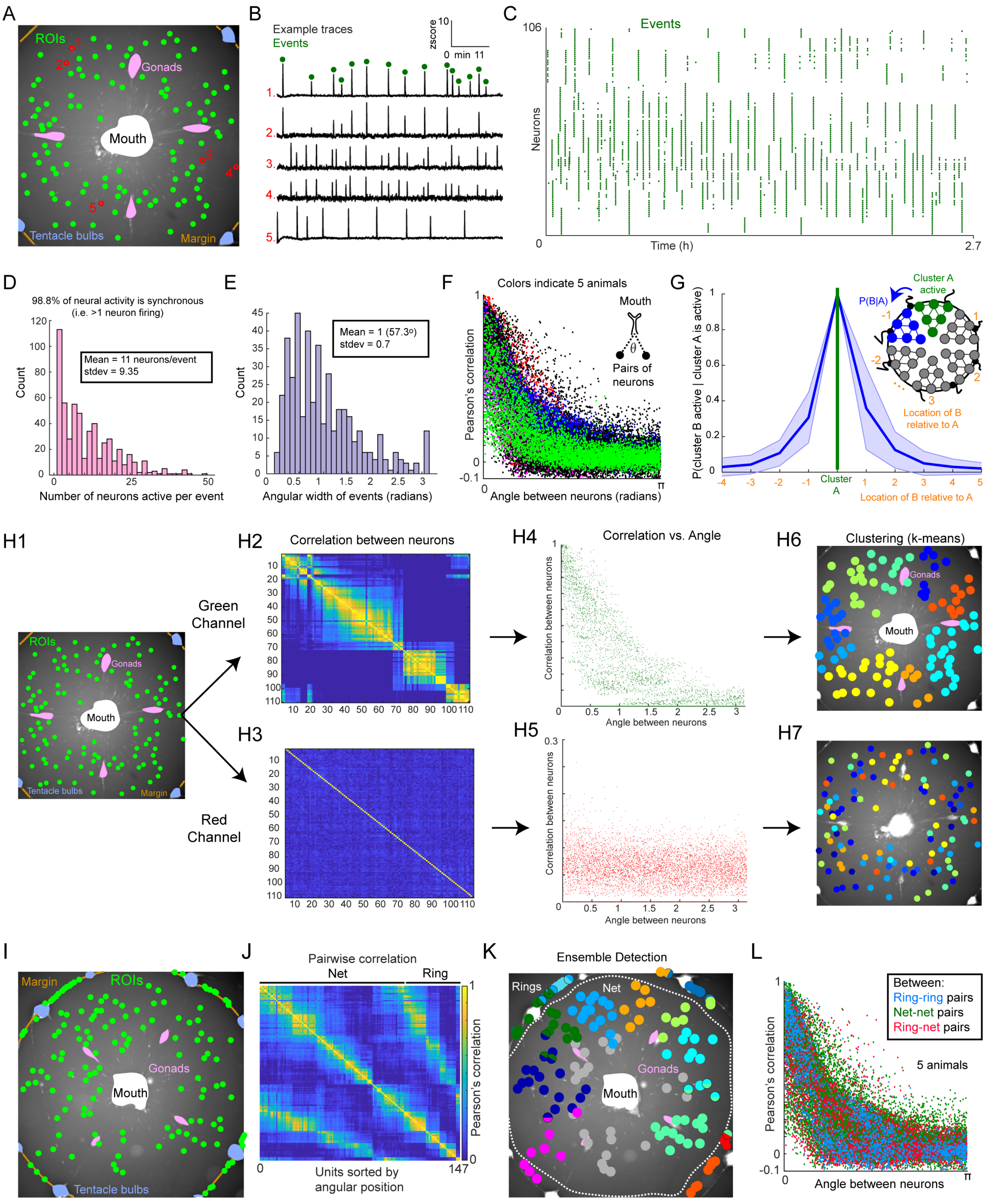

**Figure S4: RFamide neurons form radially oriented ensembles that tile the animal**

Related to Figure 5.

(A) Example field of view (FOV) for calcium imaging in a highly restrained jellyfish. Center of ROIs (green circles) shown superimposed on an image of the mCherry channel. FOV in panels A, H, I, and K is ~3.5mm$^2$. Repeated from Figure 5A with example ROIs indicated and numbered in red.

(B) Example GCaMP traces corresponding to the five red ROIs indicated in A. Green dots above the top trace show examples of "events" of neuronal activity in that trace; i.e. peaks in the GCaMP signal.

(C) Neuronal activity over time from the animal shown in panel A, with green dots indicating the GCaMP peaks for each neuron ("events"). Neurons on the y axis are sorted according to their position in the animal by angle relative to the mouth.

(D) 6065 single neuron activations form 551 single or ensemble events, pooled from recordings of spontaneous activity across the whole nerve net from 4 animals. Synchronicity (98.8%) is the number of individual neural activations (6065 total) where >1 neuron was active within a 3 second window (5990). There were 75 instances of a neuron firing alone.

(E) Angular width of events. 438 events from 4 animals; only events with >2 neurons included.

(F) Pairwise correlation of neural activity versus the angle between those neurons relative to the mouth. Data pooled from and colored by 5 individual animals.

(G) Conditional probability of a cluster "B" activating given that a cluster "A" activated. "Activation" was defined as >60% of neurons within a k-means cluster firing within a 2 second window. Comparisons were aligned such that the comparison to self is centered (P = 1), and neighboring clusters going clockwise and counter-clockwise around the animal are shown to the left and right, respectively. Blue line shows mean ± standard deviation; data pooled from n=3 animals.

(H1-7) An analysis to control for imaging or motion artifacts in which we examined fluorescence over time extracted from the red versus green channels. Pairwise correlation between fluorescence intensity over time in the red (H3) vs. green (H2), the pairwise correlation vs. angle for red (H5) vs. green (H4), and k-means clustering for red (H7) vs. green (H6) are shown. There is no similar structure in the red channel to what is observed in the green channel.

(I) FOV and ROIs for the animal shown in panels J and K, and Figure 5H-I.

(J) Pairwise correlation between neurons for the animal shown in panel I. Whether those neurons are located in the ring or net is indicated on the top.

(K) Ensemble membership using NMF/ICA for the animal shown in panel I. Neurons participating in multiple ensembles are indicated with multi-colored circles. Grey circles indicate neurons that were not assigned to an ensemble.

(L) Pairwise correlation of neural activity versus the angle between neurons relative to the mouth. Each dot is colored based on whether it was a comparison between net neurons, between ring neurons, or between a ring neuron and a net neuron. Data is pooled from 5 animals, and is the same as in panel B, but colored by comparison type rather than individual animal.

Supplemental Figure 5

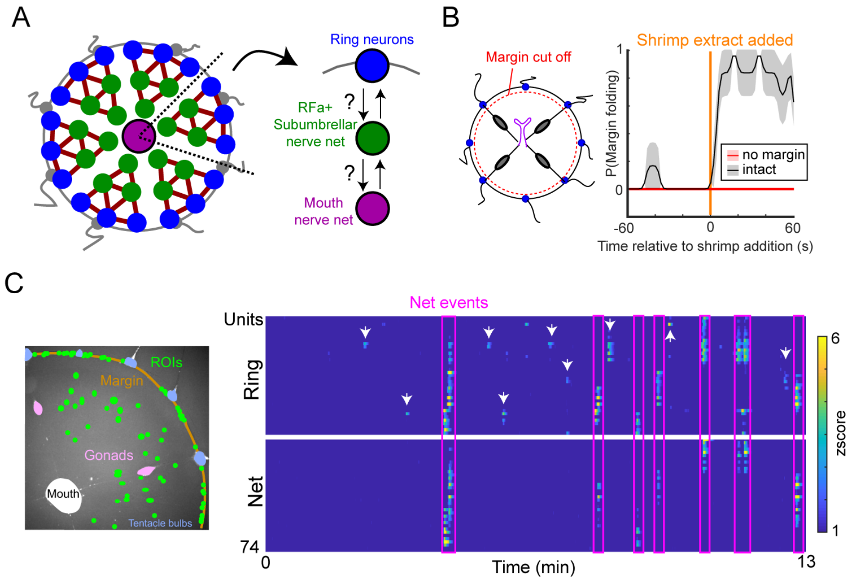

**Figure S5: The nerve rings act upstream of the nerve net**

Related to Figure 5.

(A) Schematic illustrating the questions of whether there is directionality to information flow between ring and net neurons, and whether activity travels unidirectionally or bidirectionally in the RFamide nerve net.

(B) The margin is required for folding behavior. Stimulus-triggered average from 4 animals after removal of the margin (red line). Intact control data (grey line) is the same as shown in Figure 2E, repeated here for ease of comparison.

(C) Image of FOV with ROI locations indicated (left) and activity of those neurons over time (right). Magenta boxes indicate nerve net events, and show that there is corresponding activity in the nerve rings for each of them. In contrast, white arrows point to example events in the nerve ring that do not have a nerve net correlate.

Supplemental Figure 6

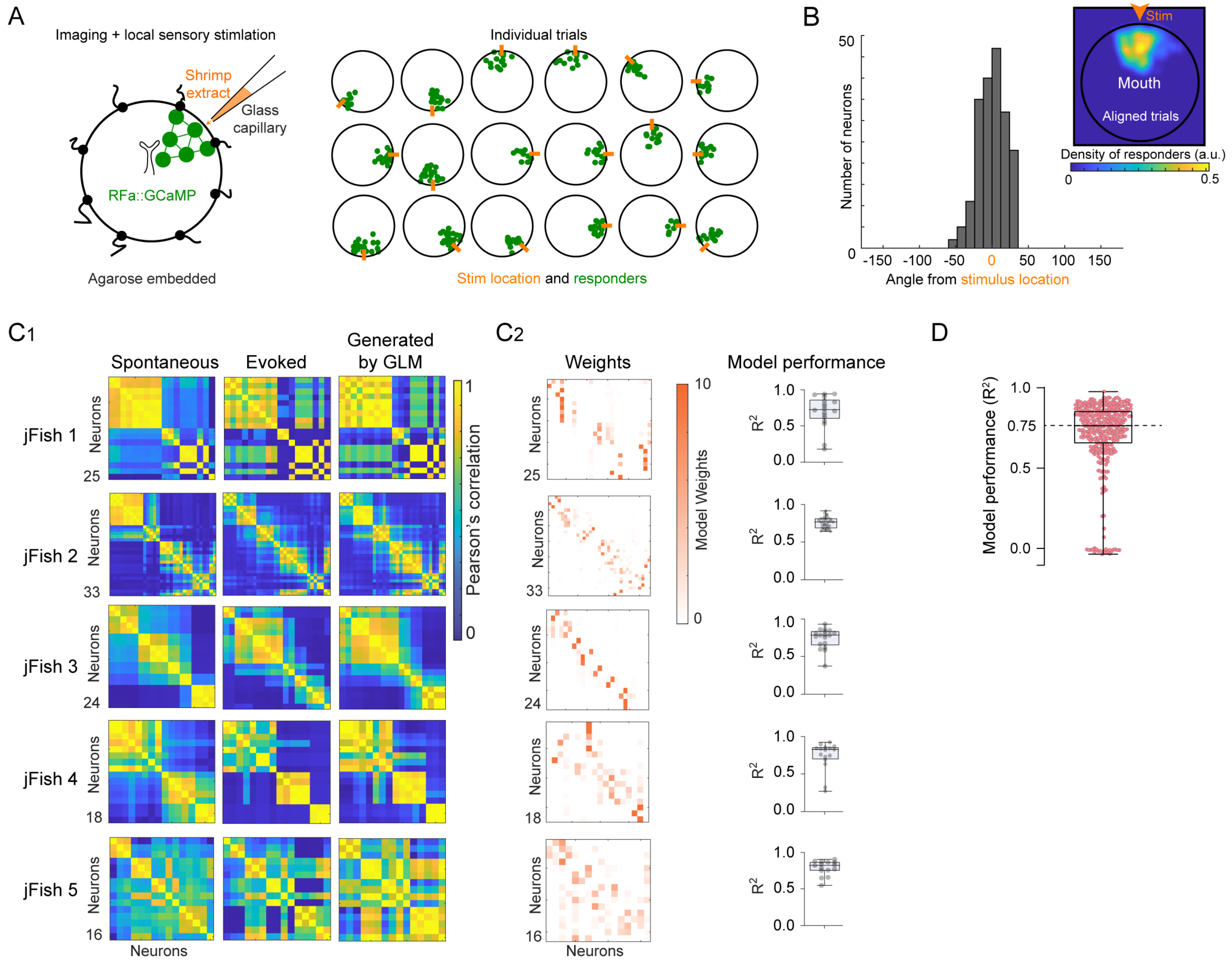

**Figure S6: Ensemble activity arises from underlying structure**

Related to Figure 6.

(A) To establish a paradigm for evoking RFamide neuron activity, we first confirmed that RFamide neurons respond upon targeted administration of food cues, which triggers spatially corresponding margin folding in unrestrained animals (Figure 2D; Supplemental Video 3). We applied droplets of food cues to the margin of agarose-embedded *Clytia* using a Femtojet and pulled glass electrodes, while performing GCaMP imaging of the nerve net. As expected, local stimulation activated a spatially localized ensemble, but did not activate distant neurons. Diagram of experiment (left), and results of individual trials (right), with the approximate location of the stimulus shown in orange and the responding neurons shown in green. Active neurons were annotated as in Figure 4E.

(B) Histogram and inset heatmap showing aligned summary data from the trials in panel A.

($C_1$) Left two panels show the correlation matrices from spontaneous versus evoked imaging epochs, repeated from Figure 6C, together for comparison with the correlation matrix generated by GLMs predicting evoked activity after training on spontaneous activity, as described in Figure 6F. ($C_2$) The weights between neurons that emerge from the GLMs after training on spontaneous activity epochs (left), and their performance on evoked data (right).

(D) GLM performance on held out data from recordings of spontaneous activity analyzed in Figure 5, which were used to generate the analyses shown in Figure 6G-I. n=339 neurons from 3 animals.
